# Tracking interlinked microbial and geochemical succession over decades in landfilled municipal solid waste

**DOI:** 10.64898/2026.02.10.705041

**Authors:** Kimber E. Munford, Daniel S. Grégoire, Laura A. Hug

**Affiliations:** Department of Biology, University of Waterloo, Waterloo, Ontario, Canada; Department of Chemistry, Carleton University, Ottawa, Ontario, Canada

**Author notes:** Address correspondence to Kimber E. Munford,.

**Keywords:** Landfill, municipal solid waste, landfill leachate, metagenomics, metals, biogeochemistry, leachate recirculation, microbial ecology

## Abstract

Landfills are heterogeneous built environments embedded in natural freshwater systems. They pose increasing risks of groundwater contamination from metal-bearing leachates over time. The interlinked succession of waste decomposition processes, microbial community membership and metal cycling across a landfill’s lifespan have not been explored, reducing our ability to predict the long-term environmental impacts of landfills. Working with 1,647 metagenome-assembled genomes from a single landfill, from samples spanning over 39 years of waste decomposition, we identified changes in landfill biogeochemistry and connected these changes to shifts in microbial community composition and predicted functions over time. Comparing between Older (aged 31 – 39 years) and Newer (aged 3 – 20 years) waste cells identified significant shifts in the availability of labile carbon, redox-associated processes, and concentrations of mobile metals – all higher in Newer cells. Newer cells were dominated by chemoorganoheterotrophs, while Older cells contained higher proportions of chemolithoautotrophs and organisms with higher metabolic versatility. Metal resistance and metal cycling genes were significantly more abundant in Older cells. Using geochemistry data from time of filling to present and microbial membership across six landfill cells of different ages, we developed a conceptual model of landfill characteristics across time. This model connects redox conditions and metal fate, highlighting leachate recirculation as a key process impacting many geochemical parameters and defining site chemistry. Our work highlights the substantial changes occurring over the stabilization phase and provides a conceptual framework for understanding this critical, final stage in a landfill’s life cycle.

**Importance:** Aging landfills pose significant risks to environmental stability and are currently poorly modeled beyond ∼20 years. Our examination of a single landfill across 39 years of waste degradation was a unique opportunity to examine the impact of time within a connected system. Our work connects geochemical data, microbial membership and predicted function, as well as physical processes (*e.g.*, leachate recirculation). Our conceptual model interlinks these facets across the lifespan of a landfill, providing an empirical data-based model of landfill aging. Previous models were extrapolated from younger waste and did not include the microbial dimension – a critical facet of the landfill ecosystem. Our model clarifies processes taking place in older wastes (30+ years), including oxygen infiltration, that have important implications for methane emission and metal mobility and fate over the longer-term.

## 1.0 Introduction

Landfills are utilized across the globe to store both residential and non-residential waste, including metals from construction waste, household refuse, and electronic waste. Landfills contribute to environmental and human health concerns over the course of their expected life spans, including long-term methane emissions, punctuated and long-term groundwater contamination, and variable potential for leaching of heavy metals or other pollutants, under fluctuating redox conditions (1, 2).

Landfill refuse is generally buried in a sequential series of cells, which are each filled over a span of several years (2, 3) and expected to stably house the refuse for decades. Once closed, landfill cells go through a predictable set of biogeochemical phases as organic matter (OM) is decomposed by microbes. In phase 1 (aerobic), OM is rapidly metabolized through aerobic processes (3, 4). This phase is brief (1-2 years) as remaining oxygen is depleted via microbial respiration, leading to anoxic conditions. In phase 2 (anaerobic acid), hydrolytic, fermentative, and acetogenic bacteria continue metabolizing OM and produce organic acids, leading to decreases in pH and an overall decrease in OM (5). Phase 2 is followed by a rapid methanogenic phase (phase 3), where methane production peaks, and a slow methanogenic phase (phase 4). Methane production slows in phase 4 as labile OM sources are depleted and methanogenesis shifts to reliance on the downstream products from other organisms’ hydrolysis of polymeric carbon sources (1, 3, 6). The duration of phases 2-4 are highly variable, and dependent on substrate conditions and climate, which impact the speed of OM breakdown (4, 6). Phase 4 can last for decades. It has been proposed that, beyond phase 4, landfills will enter a stabilization phase (phase 5). Some have defined this phase by the transition to microbial activity on more recalcitrant carbon sources (*e.g.*, humic and fulvic acids) (4, 6–8). At this point, due to reduced microbial demand, oxygen entering landfill cells via passive diffusion can lead to increased oxygen availability and more oxidizing conditions.

Shifts in redox conditions over the landfill life cycle are expected to impact microbe-metal interactions and metal mobility as landfills age (3, 4, 9). Despite this, few have investigated the differences in biogeochemical conditions between older and newer landfill cells, particularly with respect to metal resistance and metal cycling (10, 11). Overwhelmingly, existing studies compare different landfills rather than cells within one landfill, which may impact comparisons of underlying chemistry. Additionally, they typically focus on carbon cycling and greenhouse gas production, omitting metals. Studies have found that the early stages of the landfill life cycle are characterized by rapid (over 5-10 years) decreases in labile organic substrates (5, 6, 12). Others have shown that landfills experience a shift from microbial communities associated with the breakdown of OM to those dominated by lithoautotrophs over time (10, 13). What is less understood is how this microbial community succession impacts metal cycling and leaching. Some studies report higher metal concentrations in newer landfills, while others predict that metals may be mobilized in the later phases of the landfill life cycle (3, 10, 11). Increasingly oxidizing conditions in phase 5 can lead to oxidation of reduced sulfides, releasing metals and reducing pH (3, 14). Additionally, organic substrates (*e.g.,* humic or fulvic substances) remaining in waste during the stabilization phase (phase 5) can form soluble complexes or colloids with metals, increasing metal mobility (3, 4).

Here we investigate microbial community structure and associated functional genes between Older (aged 31 – 39 years) and Newer (aged 3 – 20 years) landfill cells from a landfill in the Northeastern U.S. We connect these examinations of metabolic guilds and microbe-metal interactions with changes in leachate chemistry collected over more than three decades. A previous study by Grégoire *et al.* (2023) looked at microbiological and geochemical evidence for methane cycling in the same system, where their highlighted oxygen infiltration into older landfill cells. As a result, our study focused on differences in microbial communities and leachate geochemistry over time, focusing on major microbial metabolic processes involved in and controlled by redox cycling. We hypothesized that older landfill cells would host microbial communities whose functional genes had a higher potential for utilization of inorganic electron acceptors and cycling of metals while newer landfill cells would have microbial communities with functional genes more strongly associated with anaerobic fermentative processes.

## 2.0 Results

### 2.1 Site description and sampling

The sampled site is a municipal landfill located in the northeastern United States (NEUS), site anonymity requested by site engineers. The site is composed of six landfill cells (A, B, C, D, E, and F), two of which are divided into sub-cells (D into D1 and D2, and F into F1 and F2), each equipped with leachate collection systems. Landfill cells were broadly separated by age into “Older” (cells A, B, and C; aged 31 – 39 years), “Intermediate” (cells D1 and D2; aged 24 – 26 years), and “Newer” (cells E, F1, and F2, 3 – 20 years) based on their predicted landfill phase (Older = phase 4-5, Newer = phase 2-3). Landfill phases were determined by Grégoire *et al.* 2023 (1) in their work investigating methane cycling at the site, utilizing measured biological oxygen demand (BOD) and chemical oxygen demand (COD) to assign phases. Leachate samples were collected in 2019 for sequencing and metagenomic analysis (see Methods, (1)). The site owners enthusiastically provided historic and current leachate geochemistry records under a Freedom Of Information Act (FOIA) request. Notably, microbial samples come from a single point in time (sampling in 2019) whereas leachate chemistry data spans the lifetime of the cells. As a result, microbial analyses focus on comparisons between the oldest (“Older”) and youngest (“Newer”) cells. Integrations of microbial and leachate chemistry data draw on the entire dataset.

### 2.2 Taxonomic composition of microbial communities

Starting from 1,647 total bins, dereplication was performed at 99% similarity to compare shared populations across cells, yielding 1,178 metagenome-assembled genomes (MAGs) across all eight metagenomes (all cells and sub-cells). Fewer total MAGs (>70% completion, <10% contamination) were detected in Older landfill cells (138 ± 48) than Intermediate (245 ± 34.7) or Newer cells (248 ± 42.7), with 93 MAGs in cell A, 188 in cell B, 134 in cell C, 220 in D1, 269 in D2, 210 in E, 294 in F1, and 239 in F2. Only 32 dereplicated MAGs were shared among all three age groups, and only 5 dereplicated MAGs were shared between Older and Newer cells to the exclusion of Intermediate cells (Figure 1c). Beyond limited shared MAGs, there were large taxonomic differences between communities from Older and Newer cells, based on taxonomic assignment by GTDB-tk (release 226) (15) and from a phylogenomic tree encompassing MAGs from all samples (Figure 1a). Taxonomic assignment, trees, and all subsequent analyses were performed using the set of 1,647 non-dereplicated MAGs to capture cell-by-cell differences (see Methods). Of the 1,647 MAGs in the dataset, only 77 had too few hits to the required marker genes to be included in the tree. Removed MAGs, on average, had a relative abundance of <0.5%. Most of these 77 MAGs (63.6%) were within the Halobacteriota (22 MAGs) or placed within groups with small genomes that may be missing key markers (Patescibacteriota [18 MAGs], Nanobdellota [7 MAGs], Micrarchaeota [2 MAGs]). A full table of taxonomic classifications, relative abundance, tree membership, completeness and contamination, and dereplication cluster membership is available on the Open Science Framework (OSF) (https://doi.org/10.17605/OSF.IO/MSDB8)(16).

**Figure 1:**
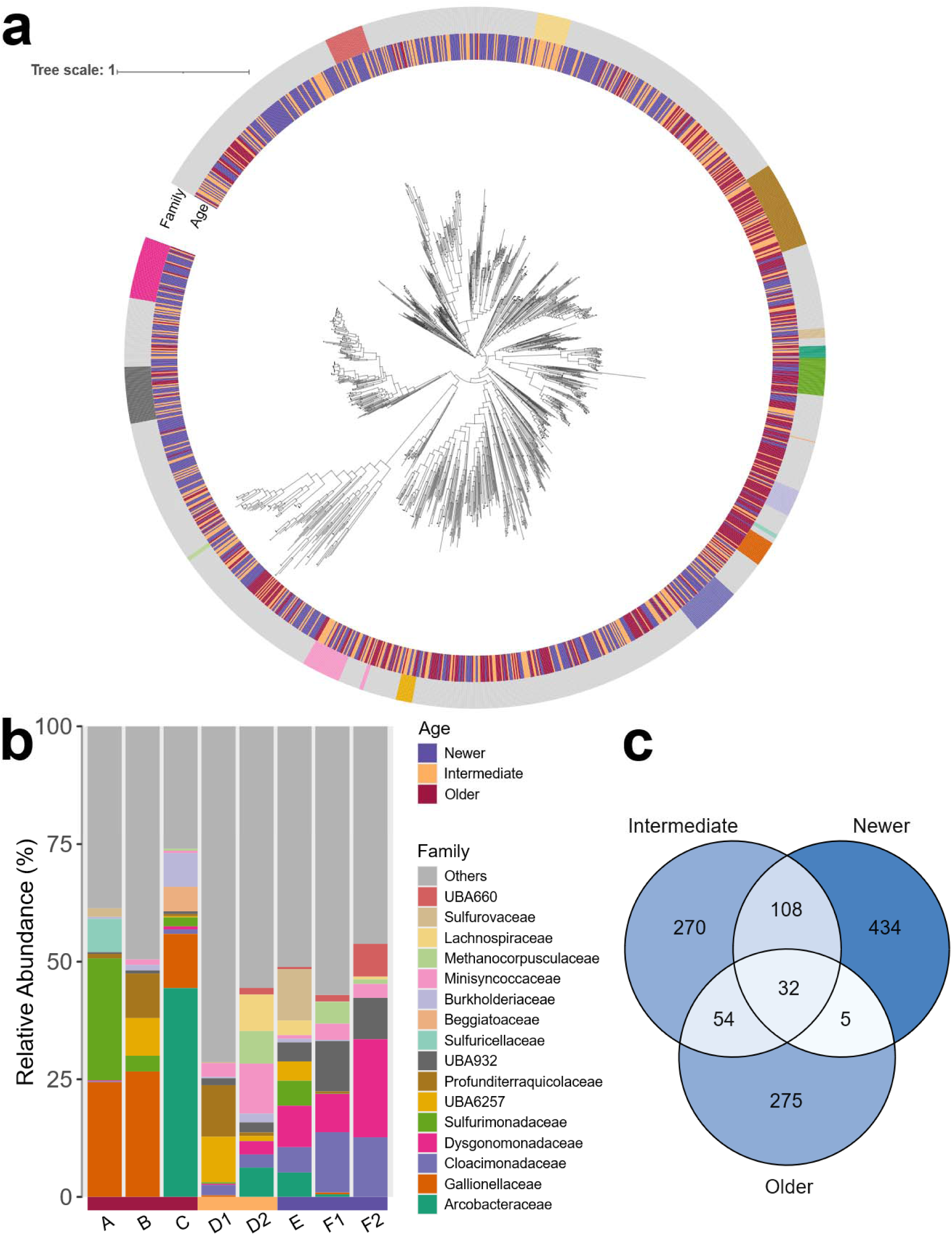
a) Phylogenomic tree based on 25 target gene HMMs for bacteria and archaea, placing 1,570 out of 1,647 MAGs in relation to each other, generated in GToTree v1.6 (58). Rings depict, outer to inner: 1) family-level taxonomy, 2) cell age category. MAGs in the tree were not dereplicated so that sample-specific relative abundances could be depicted. Colours for families correspond to those in the b) barplot of relative abundance of all bacterial and archaeal families with over 5% relative abundance in at least one landfill cell, with lower abundance lineages grouped under “Other”. Taxonomy was taken from GTDB-tk r266, where lineages starting with UBA are officially recognized but not yet formally named. c) Venn diagram indicating counts and percentages of shared MAGs (clustered at 99% similarity using DRep) among age categories across landfill cells. Venn sections are colored by proportion of MAGs within a category, with higher values corresponding to darker colors.

Overall, Older cells had lower Shannon Diversity (3.70 ± 0.640) than Intermediate (4.94 ± 0.0322) or Newer (4.63 ± 0.0445) cells. Older cells had a strong enrichment in members of the families Gallionellaceae and Sulfurimonadaceae, with enrichment in Arcobacteraceae in cell C. Collectively, members of the Gallionellaceae, Sulfurimonadaceae, and Arcobacteraceae represented 50.3% relative abundance in cell A, 30.0% in cell B, and 57.8% in cell C (Figure 1b), highlighting the overall lower microbial diversity in Older cells. The Gallionellaceae family consisted of 15 dereplicated MAGs across one unidentified (JAQTOM01 [1 MAG]) and two identified genera (*Gallionella* [10 MAGs] and *Sideroxyarcus* [4 MAGs]), and comprised 24.4% (3 MAGs), 26.7% (7 MAGs), and 11.5% (6 MAGs) in cells A, B, and C, respectively. Members of the Gallionellaceae were also present in D1 (0.327% [2 MAGs]) and F1 (0.371% [1 MAG]), but at lower abundance than in Older cells. The family Sulfurimonadaceae was represented by 24 dereplicated MAGs across 2 genera (*Sulfuricurvum* [12 MAGs] and *Sulfurimonas* [12 MAGs]) in all cells (Figure 1b, Table S1), and comprised 25.9% (6 MAGs), 3.36% (6 MAGs), and 1.91% (7 MAGs) in cells A, B, and C, respectively. Relative abundance of members of the Sulfurimonadaceae was <1% in all Intermediate and Newer cells (1 MAG in D1, 1 in D2, 8 in E, and 1 in F1). Arcobacteraceae was the most abundant family in any one sample, comprising 44.4% of the microbial community in cell C. Within cell C, three dereplicated Arcobacteraceae MAGs were present, two from the genus *Aliarcobacter* (44.0%) and one unidentified (CAIJNA01 [0.495%]). Arcobacteraceae were not detected in cells A or B but were present at lower abundances in cells D2 (6.22% [4 MAGs]), E (5.19% [1 MAG]), and F1 (0.548% [1 MAG]).

Contrastingly, Newer cells showed enrichments in members of the families Dysgonomonadaceae and Cloacimonadaceae. Dysgonomonadaceae consisted of 17 dereplicated MAGs across 2 identified genera (*Proteiniphilum* [10 MAGs] and *Petrimonas* [2 MAGs]) and 5 MAGs unidentified at the genus level. Relative abundance of MAGs in the genus *Proteiniphilum* was 3.04% (5 MAGs), 7.45% (7 MAGs), and 17.2% (10 MAGs) in E, F1, and F2, respectively, while it was below 1% in Older cells (3 MAGs in C only) and D1 (1 MAG), but slightly more abundant in D2 (2.32% [3 MAGs]). Relative abundance of MAGs in the genus *Petrimonas* was 2.12% (5 MAGs), 0.392% (7 MAGs), and 2.00% (10 MAGs) in E, F1, and F2, respectively, and less than 1% in Older (1 MAG in A and 3 in C) and Intermediate (1 MAG in D1 and 3 in D2) cells. In total, relative abundance of all unidentified members of the Dysgonomonadaceae family were 3.66% (3 MAGs), 0.319% (2 MAGs), and 1.61% (4 MAGs) in E, F1, and F2, respectively, and they were only detected in one other cell (D2 at 0.393% [2 MAGs]). The family Cloacimonadaceae consisted of 18 dereplicated MAGs, most of which were unidentified at the genus level (UBA5456 [10 MAGs]), with the identified genera being *Syntrophosphaera* (4 MAGs) and *Cloacimonas* (4 MAGs). Relative abundance of members of the Cloacimonadaceae was 5.38% (10 MAGs), 12.8% (9 MAGs), and 12.7% (6 MAGs) in E, F1, and F2, respectively. Cloacimonadaceae abundances were lower in Intermediate cells, with 2.21% in D1 (3 MAGs) and 2.80% in D2 (6 MAGs), and at less than 1% in all Older cells (1 MAG in A and 2 in C).

As shown above, Intermediate samples tended to be variable, with microbial composition in D1 more strongly resembling Older cells, while composition in D2 was closer to Newer cells. Interestingly, both subcells of cell D were enriched in families whose members are typically very small cells (≤0.8 μm) with reduced genomes (17, 18). This included the family Profunditerraquicolaceae (D1: 10.9%, D2: 0.749%) with 43 MAGs from 2 identified genera (*Undivivens* [D1: 0.449%, D2: 0.242%] and *Sherwoodlollariibacterium* [D1:0.353%, D2: 0.241%]), each represented by 1 dereplicated MAG, and 41 MAGs from unidentified genera (D1: 10.14% [24 MAGs], D2: 0.266% [2 MAGs]). Members of the Profunditerraquicolaceae were also found in Older (30 MAGs in A, B, and C) and Newer (2 MAGs in F1 only) cells at less than 1% relative abundance in all cells except B, where relative abundance was 9.47% (22 MAGs).

Similarly, two families within the Patescibacteriota phylum were also enriched in the Intermediate (D) cells. Members of the Minisyncoccaceae, represented by 17 MAGs across 2 identified genera (*Microsyncoccus* [3 MAGs] and *Minisyncoccus* [2 MAGs]) and 4 unidentified genera (12 MAGs), had 2.94% (5 MAGs) and 10.6% (8 MAGs) relative abundance in D1 and D2, respectively. Members of Minisyncoccaceae were also present in Newer cells, with 0.762% (1 MAG), 3.44% (6 MAGs), and 2.97% (5 MAGs) relative abundance in E, F1, and F2, respectively. Relative abundance of Minisyncoccaceae was less than 2% in all Older cells, detected only in B and C with 3 MAGs in each cell. Finally, an uncultured order (UBA6257) in the Patescibacteriota phylum (18 MAGs) had high relative abundance in cell D, with 7 MAGs in D1 and 3 MAGs in D2, accounting for 9.61% and 1.04% relative abundance, respectively. MAGs from UBA6257 were also present in Older (B: 8.75% [4 MAGs], C: 0.248% [1 MAG]) and Newer (E: 5.45% [7 MAGs], F1: 0.723% [1 MAG], F2: 0.178% [2 MAGs]) cells.

Although there were some similarities at the Family level, few dereplicated MAGs (5 total) were shared between Older and Newer cells (Figure 1c). In contrast, 42% of dereplicated MAGs in Intermediate cells were shared with Newer, Older, or both. Thus, Intermediate cells reflect a transitionary state between Older and Newer, highlighting how few taxa can persist following transition between landfill phases. Due to the clear distinction between microbial community compositions in Older and Newer cells, we opted to focus on comparisons between these two age groups for our downstream analyses.

### 2.3 Microbial diversity and functional genes

There were substantial differences in predicted functional capacity for microbial communities in landfill cells of different ages. In total, we examined 200 different functional gene groups and pathways. We utilized Distilled and Refined Annotation of Metabolism (DRAM) v1.0 (19) to annotate MAGs, focusing on ninety-three functional gene groups and pathways, covering central metabolism, carbohydrate-active enzymes (CAZymes), conversion of short-chain fatty acids (SCFAs) and alcohols, methanogenesis and methanotrophy, nitrogen cycling, and sulfur cycling. Annotations from four functional groups were identified using FeGenie (20), focusing on iron cycling. Ten more were annotated with an As metabolism and resistance pipeline from Dunivin *et al.* (21), and 93 were derived from BacMetScan (22), covering microbial metal interactions (respiration and resistance). Genes which contributed to functional gene categories across the different annotation tools are listed in Table S2. Only genes which were significantly different at p < 0.05 are reported here.

To investigate differences in functional potential between cells of differing ages, we integrated all annotations into a presence-absence table for all functional genes, groups, or pathways (available on the Open Science Framework at https://doi.org/10.17605/OSF.IO/MSDB8) (16). DRAM estimates coverage of metabolic pathways and completion of electron transport chain (ETC) complexes based on the percentage of genes recovered. For our presence-absence assessment, we considered pathways and ETC complexes to be present when DRAM’s estimate was >70% for a given MAG. Then, abundance of functional genes was calculated by summing the relative coverage of all MAGs that possessed a given functional gene, group, or pathway within a cell, representing the proportion of the microbial community with the encoded potential for a given function.

Across the full dataset of 1,647 MAGs, only 6 MAGs (from 0.107 to 0.730% relative abundance) had no functional gene hits with the 70% completion cutoff, including for central metabolic pathways. Four of these MAGs were members of the Patescibacteriota phylum, and two were members of the archaeal phylum Nanobdellota (formerly “Nanoarchaeota”). Members of both of these groups live as symbionts on prokaryotic hosts, lacking portions of some pathways essential for independent life (18, 23, 24). Genomes for these organisms ranged from 70.8 – 78.5% completion, so it is also possible that core metabolic genes were absent from our annotations due to incomplete genomes.

We investigated differences in functional gene and pathway abundance and prevalence between Older and Newer cells using Wilcoxon rank sum tests (p < 0.05) (Table S2). To explore the fermentative capacity of the microbial communities, we investigated the presence or absence of Complexes I - V of the electron transport chain and DRAM annotations associated with the breakdown of SCFAs and carbohydrates (CAZymes).

Complex IV of the ETC is associated with transfer of electrons to oxygen, the final electron acceptor in aerobic respiration, so a lack of this complex indicates likely anaerobic respiration. The absence of Complexes I - IV of the ETC indicates that an organism is likely an obligate fermenter. Overall, we found that 7 out of 8 Complexes I - IV of the electron transport chain were significantly more abundant in Older cells, including both low and high affinity oxidases, indicating that variable oxygen availability in the system supports aerobic metabolisms (Table S2, Figure S1). The relative lack of Complexes I - IV in Newer cells points to fermentative metabolisms and anaerobic respiration as dominant in Newer cells. ATPases also point to differences in metabolic processes between cell ages. F-type ATPases, which were significantly more abundant in Older cells, often couple to an ETC for oxidative phosphorylation, pointing to aerobic respiration when Complexes I - IV are also present. In Newer cells, V/A-type ATPases were significantly more abundant, possibly reflecting maintenance of proton gradients in microbes with low respiratory activity.

Functional gene groups associated with central carbon metabolism and organic matter decomposition also indicated that fermentation was the key metabolic process in Newer cells compared to aerobic processes in Older cells (Figure 2). Older cells had significantly higher abundance of genes or gene groups associated with the reductive pentose phosphate (Calvin) cycle (Older: 59.5 ± 17.7%; Newer: 19.3 ± 8.47%), reductive citrate (Arnon-Buchanon) cycle (Older: 43.1 ± 10.9%; Newer: 22.5 ± 7.99%), glyoxylate cycle (Older: 14.2 ± 19.4%; Newer: 2.12 ± 3.46%), and citrate (Krebs) cycle (Older: 60.3 ± 22.4%; Newer: 15.4 ± 14.7%), suggesting importance of both aerobic and anaerobic respiration, as well as carbon fixation in Older cells (Figure 2). Notably, across all samples, genes for the reductive acetyl-CoA (Wood-Ljungdahl) pathway, associated with anaerobic carbon fixation, were very uncommon, at 1.09 ± 1.36% in Older cells and 2.75 ± 1.02% in Newer cells.

**Figure 2:**
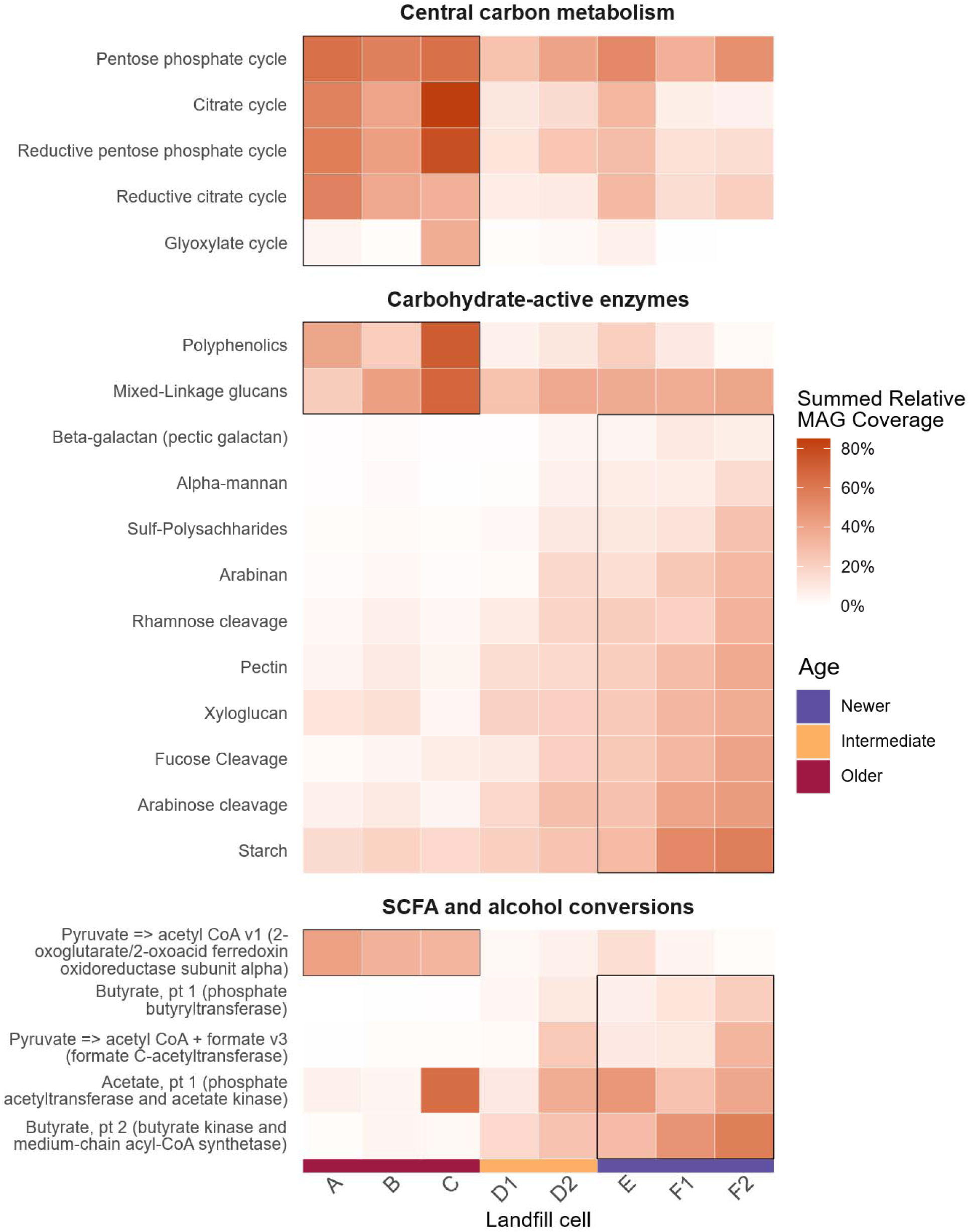
Carbon cycling differential abundance across landfill cells. Heatmap depicting the summed relative coverage of all MAGs in each landfill cell that contain genes associated with specific functions within carbon cycling. Only genes/pathways/processes which had significantly different abundances between Older and Newer cells (Wilcoxon rank sum test, Benjamini-Hochberg-corrected p-value of p < 0.05) are displayed. Boxes indicate whether abundance was significantly higher in Newer or Older cells. Central C metabolism pathways from DRAM were required to be >70% complete within a MAG for inclusion. Carbohydrate-active enzymes from DRAM either indicate the presence of genes for both backbone and oligomer cleavage (alpha-mannan, arabinan, beta-galactan, mixed-linkage glucans, pectin, starch, sulf-polysaccharides, xyloglucan) or only oligomer cleavage (arabinose cleavage, fucose cleavage, rhamnose cleavage). For polyphenolics, multiple oxidative genes are accounted for. For DRAM SCFA and alcohol conversions, the gene(s) of interest are listed in brackets. Further information can be found in the DRAM documentation.

Contrastingly, Newer cells had significantly higher abundance of gene groups associated with breakdown of carbohydrates. These included CAZymes for breaking down complex carbohydrates like pectin, starch, mannans, and xyloglucan, with 10/16 identified CAZymes at significantly higher proportions in Newer cells. The only exceptions were CAZymes for polyphenolics and mixed-linkage glucans, which were at significantly higher abundance in Older cells. The relatively higher abundance of CAZy genes for polyphenolic and mixed-linkage glucans in Older cells may point to more complex carbon sources remaining in Older cells. Additionally, Newer cells had higher abundance of some genes and gene groups associated with the breakdown of SCFAs and alcohols (5 out of 11 genes were significantly different), such as acetate, butyrate, and pyruvate (Figure 2). Enrichment in CAZy genes and genes associated with SCFA breakdown further point to fermentative activity in Newer cells.

Finally, there were also significant differences in genes and gene groups associated with methanogenesis and methanotrophy between Older and Newer cells (Table S2; Figure S2). Briefly, genes associated with methane oxidation were significantly more abundant in Older cells, further indicating possible oxygen infiltration alongside remaining methane in Older cells, though overall relative coverage of MAGs with those genes was low (<2%) (Figure S2). Trends in methanogenesis and methanotrophy were explored in further detail in Grégoire *et al.*, 2023 (1).

Older cells had higher abundance of MAGs predicted to utilize a variety of alternative electron donors and acceptors. Based on DRAM annotations (for nitrogen [N] and sulfur [S]) and FeGenie results (for iron [Fe]), we found that functional genes and gene groups for reduction and oxidation of all three elements were significantly more abundant in Older cells, pointing to metabolic flexibility in Older cells (Figure 3). Notably, genes associated with all steps of denitrification (nitrate -> nitrite [*napA* or *narG* gene] -> nitric oxide [respiratory nitrite reductase gene] -> nitrous oxide [nitric oxide reductase gene] -> nitrogen [nitrous-oxide reductase gene]) were significantly more abundant in Older cells, with abundance of these genes ranging from 11.5-55.9% in Older cells and from 2.63-15.5% in Newer cells (Figure 3). Similarly, abundance of the functional gene groups associated with oxidation of nitrite to nitrate (*nxrA* and *nxrB* genes) (Older: 25.2 ± 26.9%; Newer: 15.5 ± 5.1%) and bacterial/archaeal ammonia oxidation (*amoA* gene) (Older: 1.96 ± 1.71%; Newer: 0.340 ± 0.588%) were also significantly higher in Older cells. One exception was dissimilatory nitrate reduction to ammonia (*nrfA* gene), which was significantly more abundant in Newer cells (Older: 1.96 ± 1.76%; Newer: 19.3 ± 7.47%). Finally, abundance of the nitrogen fixation gene *nifH* was significantly higher in Older cells (Older: 38.7 ± 21.4%; Newer: 7.61 ± 4.13%). Taken together, the primary processes for generating biologically available nitrogen differ between cell age categories. There was evidence for both aerobic and anaerobic processes, including N fixation, in Older cells, whereas genes for dissimilatory nitrate reduction to ammonia, a process common in strongly reducing environments, were more abundant in Newer cells. These results highlight the differing mechanisms of N acquisition for landfill microbiota as landfills age.

**Figure 3:**
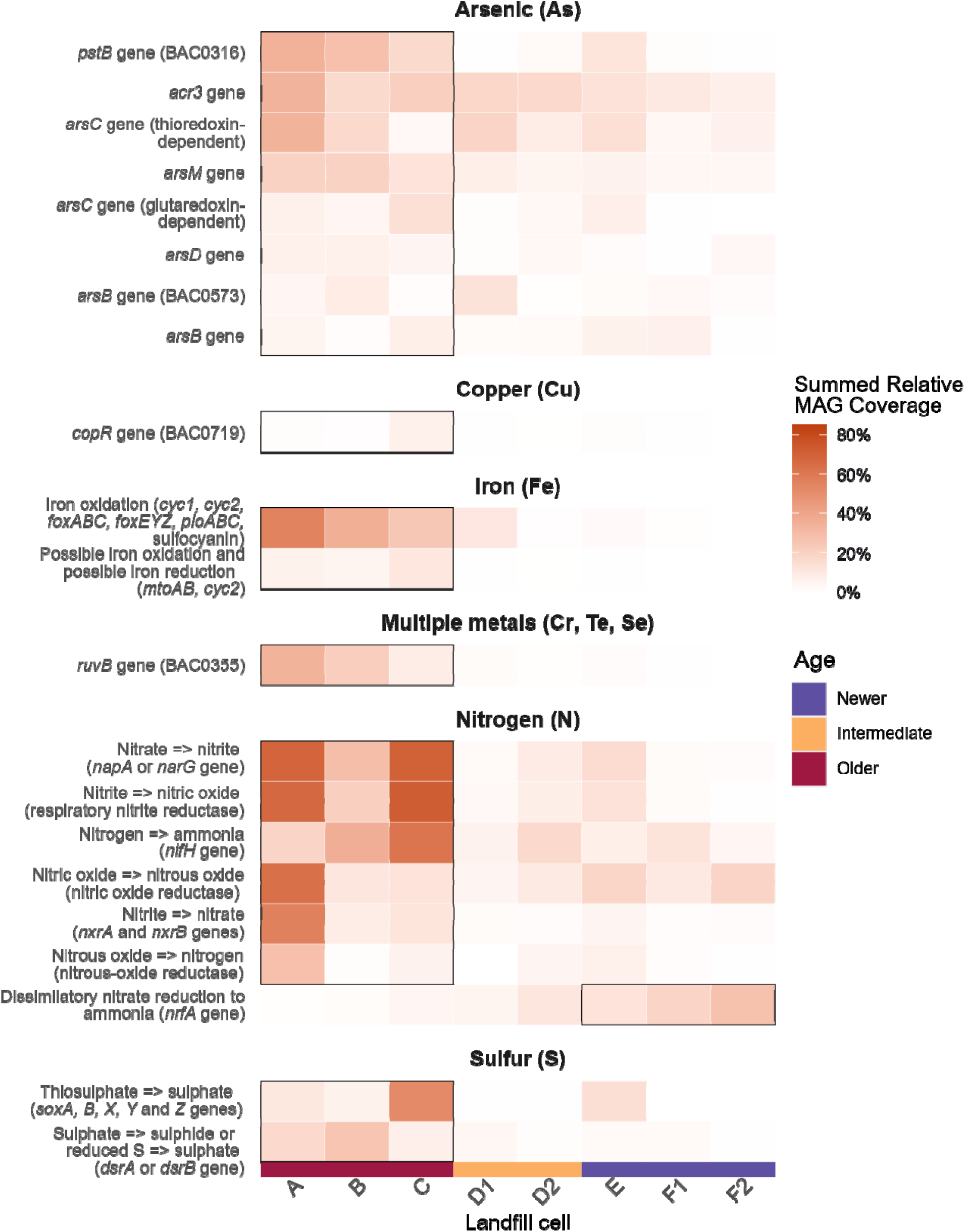
Redox cycling and metal resistance differential abundance across landfill cells. Heatmap depicting the summed relative coverage of all MAGs in each cell that contain genes/processes associated with specific functions within redox cycling and metal resistance. Only genes/processes which had significantly different abundances between Older and Newer cells (Wilcoxon rank sum test, Benjamini-Hochberg-corrected p-value of p < 0.05) are displayed. Genes/processes with a maximum of < 0.05% relative coverage were pruned for clarity and are presented in Figure S3. Genes and functional groups come from four tools: FeGenie and an HMM set from Dunivin et al. for Fe and As, respectively, BacMetScan for metal-resistance genes and DRAM for functional groups associated with S and N cycling. For As-related genes detected using the Dunivin et al. HMM set, the gene of interest is listed. For BacMetScan annotations, the BacMet number is listed in brackets following the gene. For FeGenie annotations, the process is listed, followed by the relevant genes in brackets. Finally, for DRAM annotations, the process is listed, followed by the gene(s) of interest in brackets.

Functional gene groups for reduction and oxidation of S and Fe followed similar trends. Two functional gene groups for the oxidation or reduction of sulfate were significantly different in abundance between Older and Newer cells. The functional gene group for partial or complete oxidation of thiosulfate to sulfate (including *soxA, soxB, soxX, soxY* and *soxZ*) was highly abundant in Older (22.7 ± 25.5%) cells, in contrast to Newer (4.70 ± 8.14%) cells. Interestingly, the functional gene group for dissimilatory reduction or oxidation of sulfate (*dsrA* or *dsrB*) was also much higher in Older (16.5 ± 9.33%) than Newer (1.84 ± 1.35%) cells (Figure 3).

Functional gene groups identified by FeGenie as associated with iron oxidation (Older: 38.5 ± 15.5%; Newer: 0.875 ± 1.00%) and possible iron oxidation or reduction (Older: 7.09 ± 2.92%; Newer: 0.890 ± 0.154%) were also significantly more abundant in Older cells (Figure 3; Table S2). The increase in sulfate and iron reduction genes in Older cells relative to nitrate reduction genes in Newer cells may reflect shifts in terminal electron acceptors as substrates are depleted over time. Finally, abundance of arsenic metabolism genes (*e.g.*, *arrA*, *aioA*) were not significantly different between Older and Newer cells across either the Dunivin *et al*. HMM set (21) or via the BacMet database. The heightened abundance of genes or gene groups for both oxidation and reduction of Fe, S, and N compounds in Older cells highlights a greater reliance on alternative electron donors and acceptors in Older cells.

Although there were no significant differences in abundance of respiratory arsenic metabolism genes, genes associated with As detoxification and tolerance were significantly different. Abundance of the arsenic methyltransferase *arsM* (Older: 17.3 ± 4.30%; Newer: 4.30 ± 1.23%), and two cytoplasmic arsenate reductase genes (thioredoxin-dependent and glutaredoxin-dependent) were also significantly higher abundance in Older cells. Thioredoxin-dependent arsenate reductase was more abundant in general (Older: 17.8 ± 15.3%; Newer: 7.80 ± 5.46%) than glutaredoxin-dependent arsenate reductase (Older: 8.06% ± 5.27%; Newer: 2.41 ± 3.88%). Both are associated with cytoplasmic reduction of As(V) to As(III) so it can be pumped out of the cell via an arsenite efflux pump (*acr3* or *arsB*). Both arsenite efflux pump genes were significantly more abundant in Older cells, with arsenical resistance-3 protein (*acr3*) (Older: 23.6 ± 8.96%; Newer: 9.78 ± 2.95%) being more abundant than *arsB*, which was detected through both the Dunivin *et al*. HMM set and BacMetScan. The two annotation tools use different methods to annotate genes (21, 22), and there was little overlap in annotation of *arsB* between the two methods, with 70 MAGs identified as containing *arsB* from the Dunivin *et al*. set and 99 from BacMet, but only 4 overlapping between the two. Across both annotation tools, abundance of the gene encoding the arsenite efflux pump (*arsB*) ranged from 1.31 to 8.43% in Older cells and from 0.266 to 5.99% in Newer cells. The arsenic metallochaperone (*arsD*), also significantly more abundant in Older cells (Older: 5.76 ± 1.39%; Newer: 1.84 ± 1.55%), chaperones intracellular As(III) to *arsB*-encoded efflux pumps. Higher relative abundance of genes that encode various parts of the As detoxification system point to more prevalent As-resistance in Older cells relative to Newer cells. Additionally, the phosphate-transporting ATPase (*pstB*), which can bring As(V) into cells, was also significantly more abundant in Older cells (Older: 26.1 ± 9.30%; Newer: 4.36 ± 6.21%), indicating the possibility for increased uptake of As in Older cells as a consequence of P scavenging.

We also looked for other metal-resistance genes using BacMetScan and the BacMet database, finding that Older cells displayed significantly higher abundance of genes associated with mitigating chromate toxicity (*chrA*, *chrF*), copper resistance and metabolism (*copR*), cobalt-zinc-cadmium resistance (*czcA*), cobalt, cadmium, and nickel transport (*dmeF*), tellurium resistance (*terD*), and repairing DNA damage following chromate, tellurite, or selenite exposure (*ruvB*) (Figure 3; Figure S3). The DNA repair gene encoding ATP-dependent DNA helicase (*ruvB*) was the most abundant of these metal resistance genes (Older: 21.1 ± 12.8%; Newer: 0.660 ± 0.843%). All the other metal resistance genes listed above were below 3% relative coverage, and, interestingly, five of six of these (*chrA*, *chrF*, *czcA*, *dmeF*, and *terD*) were not found in any MAGs from Newer cells. The gene encoding the copper resistance regulator (*copR*) was also below 3% abundance on average but present in both Older (2.66 ± 3.10%) and Newer (0.404 ± 0.495%) cells.

### 2.4 Leachate chemistry

Leachate chemical parameters were investigated for the 5 years prior to sampling and the period of filling for individual cells. Key parameters were selected that associate with microbial functions of interest (Table 1, from full set in Table S3). Tables of all leachate chemistry parameters over time have been uploaded to the Open Science Framework (https://doi.org/10.17605/OSF.IO/MSDB8) (16). Further supporting the differential abundance of redox-cycling genes discussed above, conditions were broadly less reducing in Older cells relative to Newer cells (Table 1). BOD and COD were both significantly lower in Older cells in the 5 years prior to sampling, and there was a massive decline in both BOD (from 9629 ± 9940 to 71.6 ± 194.2 mg/L) and COD (from 14285 ± 13298 to 446 ± 387 mg/L) since the time of filling, indicating a clear decrease in labile carbon sources (Table 1; Table S3). Total organic carbon (TOC) in leachate was also significantly lower in Older cells in this time window. Other carbon sources were also at significantly lower concentrations in Older cells, including total phenolics, and acetic acid and propionic acid, both SCFAs. The other investigated SCFAs were not statistically significantly different but had overall lower concentrations in Older cells (Table S3). Shifts in BOD, COD, and labile carbon sources over time and between Older and Newer cells reflect differences in abundance of functional genes related to fermentation and carbon metabolism over time. Interestingly, leachate concentrations of metals (As, Cr, Fe, Ni) and redox-associated parameters (reduced sulfide and ammonia) were significantly higher in Newer cells than Older cells in the 5 years prior to sampling, despite lower abundance of functional genes associated with sulfate reduction and metal resistance in Newer cells (Table S3). This may be linked to sulfate reduction decreasing soluble concentrations of metals in Older cells, with sulfides incorporated into solid waste.

**Table 1:**
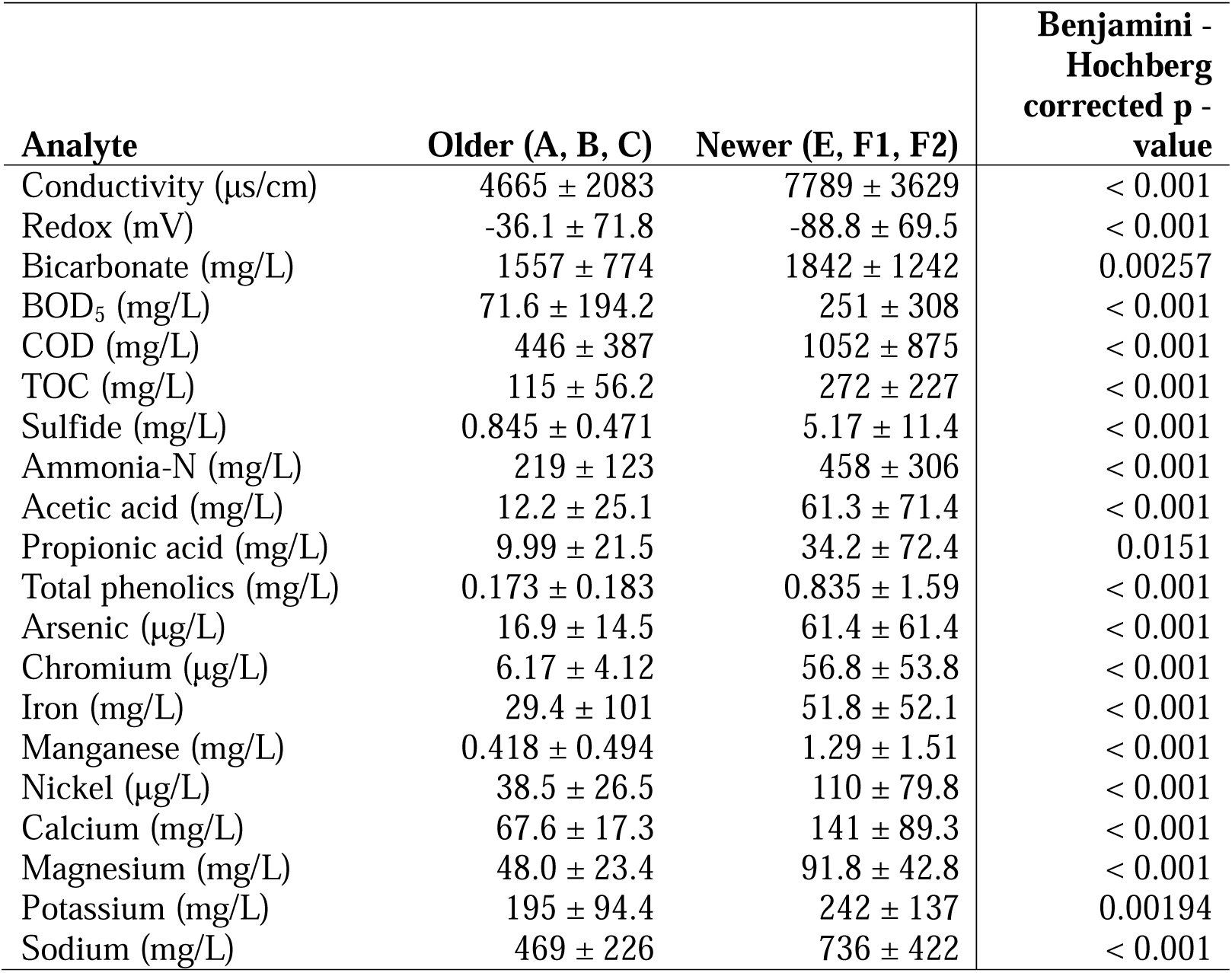
All geochemical measurements (mean±sd) for key variables, collated by landfill cell age class during filling of the cells and during the 5 years prior to sampling in winter 2019. Benjamini-Hochberg corrected p-values are presented for Wilcoxon rank sum comparisons between age classes in the 5 years prior to sampling. Only parameters that were significantly different at p < 0.05 are presented here. Remaining variables can be seen in Table S3.

The 5-year window was selected to represent the geochemical conditions that had most recently shaped the microbial communities examined. It remains possible that earlier conditions have exerted a long-term impact on community membership and functional capacity. Leachate concentrations of most metals (Cd, Cr, Co, Cu, Fe, Mn, Ni, Zn) were significantly higher (p < 0.05) in Older cells at the time of filling relative to the time of sampling (Table S3). For example, concentrations of Cr in leachate decreased from 160 ± 410 to 6.17 ± 4.12 μg/L from filling to the 5 years prior to sampling, over 25 times lower. Similarly, Fe also decreased to over 25x less, from 849 ± 2960 to 29.4 ± 101 mg/L (Table S3). Microbial communities in Older cells also had significantly higher abundance of functional genes and gene groups associated with resistance to some of these metals (Cr, Cd, Cu, Co, and Zn) and Fe oxidation and reduction, possibly pointing to a link between historic metal concentrations and present-day capacity for microbe-metal interactions.

To develop a picture of changes in overall landfill chemistry over decades, we aggregated leachate chemistry data into 5-year rolling averages. We computed z-scores for each parameter to assess differences in trends over time and clustered parameters into 8 k-means clusters based on an elbow plot of within-cluster sum of squares (Figure S4, Table 2) to combine leachate chemistry variables into clusters with similar variance over time. We separated these into those associated with key explanatory parameters (conductivity, redox, pH, OM breakdown) and remaining chemical groupings, highlighting changes over time by plotting averaged z-scores (Figure 4, Figures S5-S9). This allowed us to identify key chemical parameters driving changes in landfill conditions. A table of all z-score adjusted and averaged cluster data is available on the Open Science Framework (https://doi.org/10.17605/OSF.IO/MSDB8) (16).

**Figure 4:**
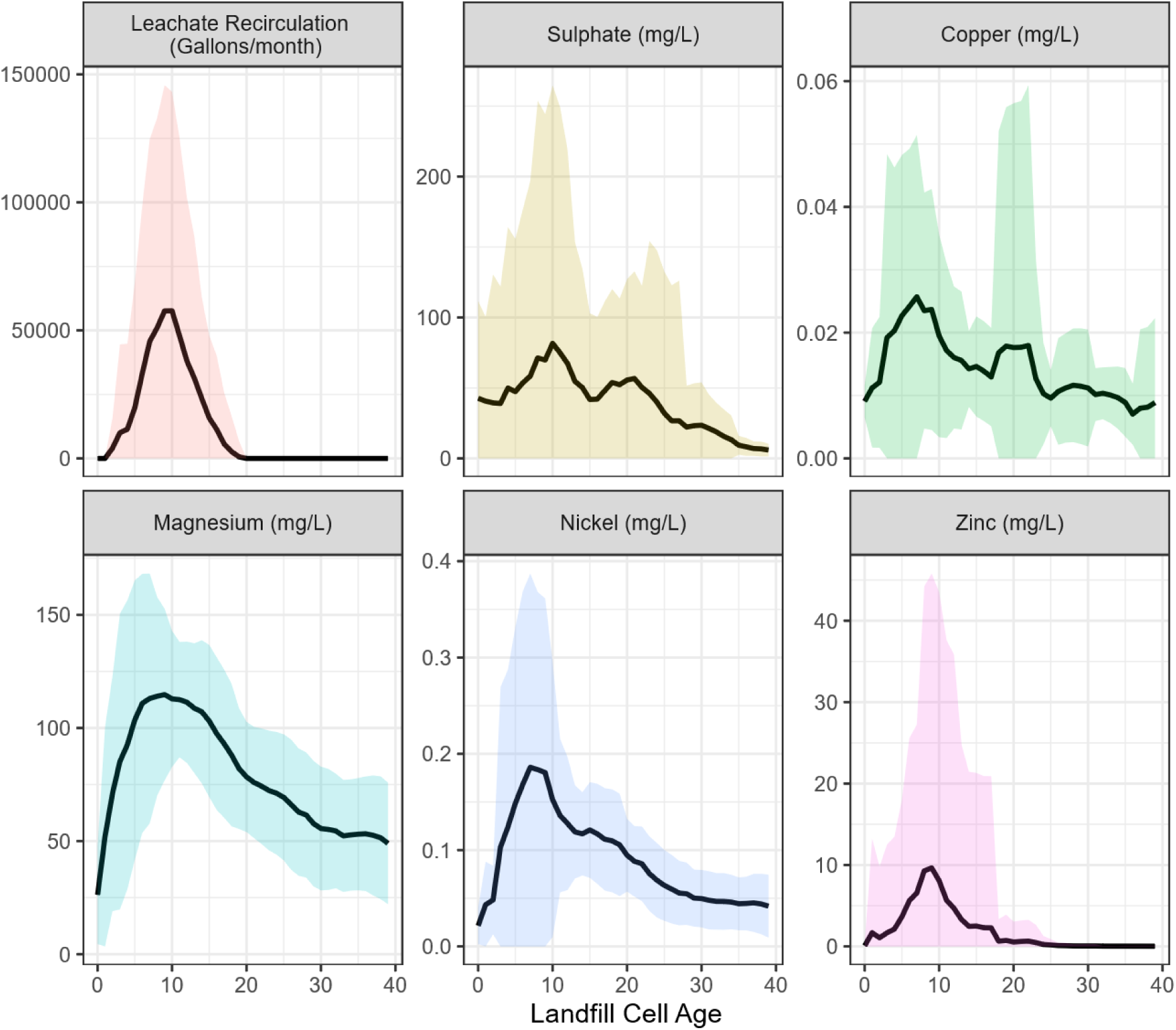
Line graphs depicting the 5-year rolling average (line) and standard deviation (shaded band around line) for landfill leachate parameters across 36 years of leachate chemistry data. All depicted parameters were within one k-means cluster (cluster 1).

**Table 2:**
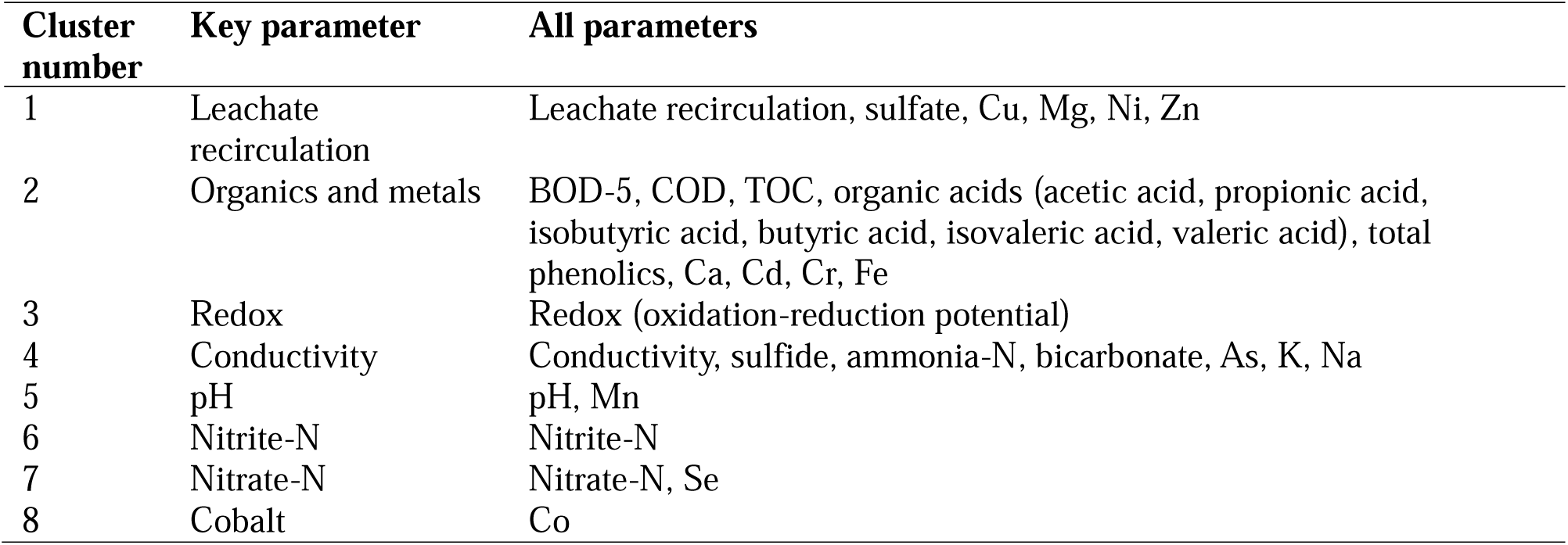
Cluster membership for 8 k-means clusters of z-score adjusted 5-year rolling averages of 36 years of landfill leachate data.

| Cluster number | Key parameter | All parameters |
| --- | --- | --- |
| 1 | Leachate recirculation | Leachate recirculation, sulfate, Cu, Mg, Ni, Zn |
| 2 | Organics and metals | BOD-5, COD, TOC, organic acids (acetic acid, propionic acid, isobutyric acid, butyric acid, isovaleric acid, valeric acid), total phenolics, Ca, Cd, Cr, Fe |
| 3 | Redox | Redox (oxidation-reduction potential) |
| 4 | Conductivity | Conductivity, sulfide, ammonia-N, bicarbonate, As, K, Na |
| 5 | pH | pH, Mn |
| 6 | Nitrite-N | Nitrite-N |
| 7 | Nitrate-N | Nitrate-N, Se |
| 8 | Cobalt | Co |

Interestingly, we found that a group of parameters (SO_4_^2-^, Cu, Mg, Ni, Zn) clustered with the historic practice of leachate recirculation at the site, co-varying over time, with all parameters peaking in concentration at around 10 years since cells began filling (Cluster 1, Figure 4). Leachate recirculation occurred off and on within the landfill from 1985 to the end of 2009. To investigate the impact of leachate recirculation as a driver of leachate chemistry, we performed simple linear regressions on z-score-scaled 5-year rolling averages for all cell ages where there was leachate recirculation (Table S4). Parameters significantly influenced (p < 0.05) by the amount of recirculated leachate included carbon-associated parameters (BOD_5_, COD), some metals (Cu, Cr, Fe, Ni, Zn), Mg, oxidation-reduction potential, and two redox-associated nutrients (nitrite-N, sulfate). Given that leachate recirculation is generally performed to maintain reducing conditions and speed up waste decomposition, increases in BOD_5_/COD and decreases in oxidation-reduction potential are to be expected. The increases in metal concentrations may be linked to faster breakdown of waste, mobilizing more metals into solution, in addition to metals being recirculated back into the system.

The largest cluster contained parameters associated with the breakdown of carbon sources (BOD-5, COD, TOC, SCFAs, total phenolics) and some metals (Ca, Cd, Cr, Fe) (Cluster 2, Figure S5). Calcium concentrations and carbon parameters all followed a similar trend, broadly peaking just prior to the 10-year mark then declining (Figure S5). For Cd, Cr, and Fe, the rise and fall were more gradual, but still present. The peak for this cluster was earlier than the peak for the recirculation-associated group (including Cu, Mg, Ni, and Zn) (Table 2; Figure 4, Figure S5). Contrastingly, oxidation-reduction potential (sole parameter in Cluster 3) displayed a broad dip at 10 years and a slow rise afterward, reflecting potential oxygen infiltration into the aging landfill (Figure S6).

Conductivity was clustered with sulfide, ammonia-N, bicarbonate, As, K, and Na (Table 2). Conductivity and concentrations of ammonia-N, bicarbonate, K, and Na all followed a similar trend with a broad peak between 10 and 20 years (Figure S7). Sulfide and As were less similar, with a bump in concentration just before the 20-year mark (Figure S7). pH only clustered with Mn, both of which peaked at the beginning of the landfill life cycle before dropping off (Figure S8).

The remaining clusters, all depicted in Figure S9, were cobalt alone, nitrite-N alone, and a cluster containing nitrate-N and selenium. Trends in these were more sporadic over time, and total concentrations tended to be low (below 5 mg/L for nitrate-N and nitrite-N and below 1 mg/L for Co and Se).

Results for redox-associated parameters can be explained in the context of redox metabolism broadly. Manganese peaked early in the landfill life cycle, followed by Fe, both of which would likely be released into landfill leachate upon reduction from oxide forms. Then, following the large peak in Fe, there was a small increase in sulfide. This redox cascade is expected, though measured sulfide concentrations were quite low (<15 mg/L).

### 3.0 Discussion

This study investigated differences in geochemistry and microbial community structure, composition and functional genes in leachate from cells in a municipal solid waste landfill across 39 years of waste deposition. Few studies have explored the chemical and biological conditions in landfill cells in the later phases of decomposition under field conditions (phases 4 and 5) (3, 13). We compared “Older” (aged 31 – 39, phase 4/5) cells and “Newer” (aged 3 – 20, phase 2/3) cells, finding significant differences in microbial community structure and composition as well as microbial functional genes, and connected these differences to the underlying leachate chemistry. To anchor this discussion, we developed a conceptual model synthesizing results from microbial community functional predictions and landfill chemistry modeled over time (Figure 5).

**Figure 5:**
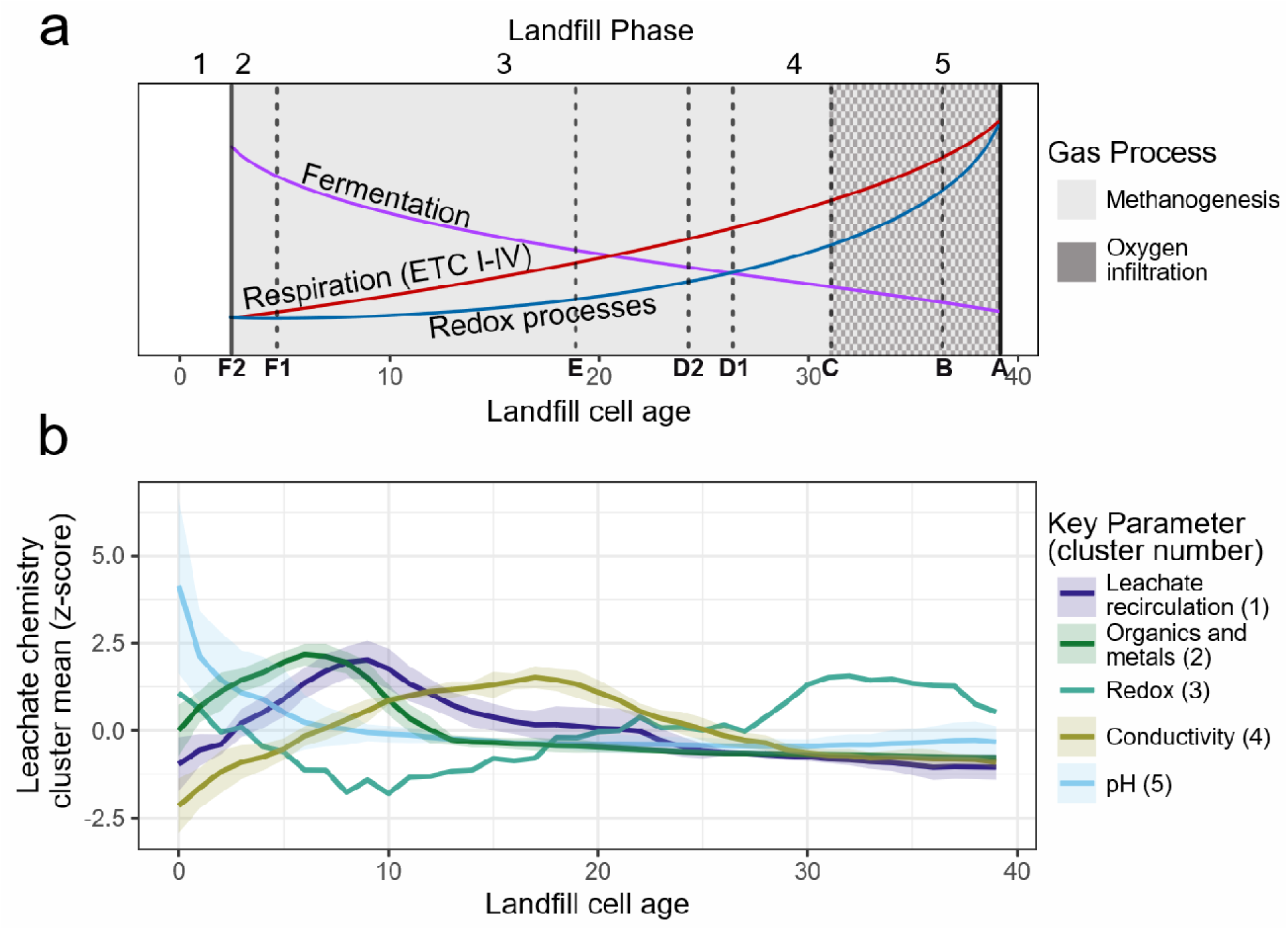
Conceptual diagram of landfill processes over time and landfill phases. a) Trends in functional gene groups over landfill cells varying in age from 3 to 39 years across 4 landfill phases, overlaid on predicted methane/O_2_ dynamics. Solid lines represent the beginning and end of our observations, and the dashed lines represent the ages at which all other cells were sampled. Landfill phases are depicted along the top, but it is notable that cells F1 and F2 (the two newest cells) were transitioning from phase 2 to 3. Trend lines represent three functional gene groups: Fermentation (SCFA/alcohol degradation, carbohydrate-active enzymes), Respiration (defined by abundance of complete (>70% of constituent genes present) electron transport chain complexes for ETC I-IV) and Redox Processes (comprising genes for redox transformations of Fe, N, S, and As). Gas dynamics are depicted as greyscale background, with methane dominating an anoxic environment from years 3-26, followed by oxygen infiltration in later years. b) Line graph depicting trends in key geochemical variables across 36 years of leachate chemistry data. Lines depict the means of all z-score-adjusted parameters within a given k-means cluster, with standard deviation of all z-score adjusted parameters within a cluster indicated by lighter bands. Clusters are named in the legend based on their key physical variable(s). Note that redox has no standard deviation because it was the sole member of its cluster (Cluster 3).

### 3.1 Microbial contributions across the 5-phase model of the landfill life cycle

This study fills important gaps in our understanding of the role microbial assemblages play in driving leachate chemical changes over the landfill life cycle, particularly in the later phases. In the early phases of the landfill life cycle, one of the key changes is an accumulation of labile organic matter (*e.g.*, organic acids) as microbes break down complex organic material (3, 12). We observed a gradual increase in leachate organic acids (acetic, propionic, butyric, isobutyric, isovaleric, and valeric acid), biological oxygen demand, and chemical oxygen demand from 0-7 years for landfill filling, before a decrease of these parameters to near zero by year 12 (Figure 5b – cluster 2). Similar trends in organic acids and BOD/COD have been found by other authors, who have observed major decreases within 10 years (12) or as little as 4 years (5). Within this first seven years, pH declined substantially due to this accumulation of organic acids, produced from hydrolysis and fermentation of OM during the anaerobic acid phase (phase 2) of the landfill life cycle (3–5). It is expected that organic acids produced in phase 2 then feed the rapid methanogenesis that characterizes phase 3, explaining the overall decline in organic acids and biological and chemical oxygen demand (4).

Our microbial samples, collected in 2019, captured two cells transitioning from phase 2 to 3 (F1 and F2) and one in phase 3 (E), making up our group of “Newer” cells. These cells had higher Shannon Diversity and higher numbers of MAGs compared to Older cells. Very few MAGs were shared between Older and Newer cells, reflecting differences in geochemical context and shifts in microbial metabolism and survival strategies as landfill waste decomposes over time (3). Due to the overall higher diversity of the microbial communities in Newer cells, only the families Dysgonomonadaceae and Cloacimonadaceae were more relatively abundant than other families across Newer cells (Figure 1b). MAGs from both families encoded a variety of genes associated with carbohydrate-active enzyme (CAZyme) families and with breakdown of SCFA and alcohol conversions. Members of the Dysgonomonadaceae have been found in a variety of environments, including landfills (25–27) and anaerobic digesters (28, 29), and are associated with the breakdown of a variety of organic compounds that are commonly present early in a landfill life cycle. In our data, the genomes of members of the Dysgonomonadaceae’s encoded genes for most of the investigated CAZymes as well as for conversions of butyrate, acetate, and pyruvate. Similarly, members of the Cloacimonadaceae have been found in landfills (30) and anaerobic digesters (31, 32), and are associated with syntrophic propionate oxidation (32). Genomes from the Cloacimonadaceae contained fewer genes for CAZymes, targeting primarily chitin and starch degradation. The Cloacimonadaceae were also predicted to be active in acetogenic metabolisms, which likely support downstream methanogenesis (1, 30).

The abundance of genes associated with fermentative processes was significantly higher overall in Newer cells. This included CAZymes for the breakdown of many polysaccharides and genes associated with SCFA and alcohol conversion (Figure 2). Further supporting the predominance of fermentative metabolism, Newer cells had a near absence of genes associated with complexes I - IV of the electron transport chain, which are necessary for respiration (Figure 5).

During phase 3 (represented by Newer cells), rapid methanogenesis is the dominant process, utilizing breakdown products from the anaerobic acid phase (*e.g.*, SCFAs). Methane was being generated in all Newer cells, with large decreases in total gas flared in Intermediate and Older cells relative to Newer cells (explored in greater detail in Grégoire *et al.* [1]). In keeping with this, Grégoire *et al.* (1) found a higher proportion of putative methanogens in Newer and Intermediate cells than in Older cells. Supporting this shift in microbial metabolism, we also observed large decreases in BOD, COD, TOC, and SCFAs, consistent with microbial consumption of labile carbon sources and a likely transition from fermentation-based processes to predominantly methanogenesis early in the landfill life cycle, corresponding to the shift from phase 2 to 3 (Figure 5).

During phase 4, total gas production tends to decrease, but methanogenesis proceeds (4, 7, 33). Reduced concentrations of SCFAs and other more labile carbon substrates by this phase lead microbes to rely on degradation of more complex OM, a slower process that reduces overall methane production. In our data, Intermediate-aged cells (D1 and D2) had more recently entered phase 4, while Older cell C was also classified to this phase. In all phase 4 cells (C and D), total gas production was substantially lower than in phases 2-3 (Newer cells) but still dominated by methanogenesis. We also observed a decrease in the abundance of genes or functional gene groups associated with fermentative functions and a corresponding increase in MAGs carrying genes that confer metabolic flexibility across a variety of redox processes.

Cell C exhibited higher abundance of functional genes and gene groups associated with redox cycling of Fe, N and S than D1/ D2, reflecting the shift toward metabolic flexibility by the end of the phase 4, as methanogenesis continues to slow (Figure 3; Figure 5). Interestingly, we also observed overall lower leachate concentrations of N, S, and Fe by phase 4, indicating a shift in microbial metabolism to lithoautotrophy as methanogenesis slowed and conditions became less reducing (Figure 5b). These changes in functional potential were in large part due to shifts in microbial community composition between cells D1/D2 and cell C. Cell C, representing a later portion of phase 4, was broadly dominated by 2 MAGs from the genus *Aliarcobacter* (44.0% relative abundance). In particular, the genomes of these MAGs encoded genes related to denitrification (nitrate > nitrite > nitric oxide), N fixation, degradation of carbon sources (polyphenolics, chitin, mixed-linkage glucans), oxidation of thiosulfate, and several genes related to acetate metabolism (acetate kinase, phosphate acetyltransferase, acetate--CoA ligase [ADP-forming] subunit alpha). The potential for metabolic flexibility, something documented across the Arcobacteraceae family (34, 35) may explain the predominance of this group in cell C. This metabolic flexibility may give them a competitive advantage as biological activity declines throughout phase 4.

Phase 5 of the landfill life cycle, called the stabilization or maturation phase by some and the humic phase by others, is characterized by an increased potential for oxygen buildup and oxidative processes, as well as organic matter that is largely recalcitrant (*e.g.*, humic and fulvic acids) (6–8). Because this phase may take decades to develop, few studies have investigated microbial communities and leachate geochemical conditions in detail. By phase 5, we observed an overall decline in carbon-associated parameters in leachate, including BOD, COD, TOC, and concentrations of SCFAs. This decrease in BOD, COD, and the BOD to COD ratio is often seen as indicative of the transition to “mature” landfill status (3, 5, 33). These declines also signify reduced overall biological activity, indicating reductions in rates of decomposition as landfill waste ages – a critical indicator of the stabilization phase. This reduction in decomposition may be linked to the observed increase in oxygen flared from Older cells at this site, indicating that a lower proportion of infiltrating oxygen is being consumed by microbes (1). This observed increase in oxygen, though still at low levels, has been proposed by several authors as an additional indicator of the stabilization phase (3, 4). We also observed an increase in oxidation-reduction potential across in this window, a further indicator of a shift toward more oxidizing conditions and processes in the later phases of the landfill life cycle.

Reflecting the decrease in biological oxygen demand, we observed fewer overall MAGs and lower diversity in Older cells (A, B, Cs) relative to Newer cells. Microbial communities in Older cells were relatively enriched in chemolithoautotrophs (*e.g.,* Gallionellaceae, Sulfurimonadaceae), contrasting with the variety of diverse putative heterotrophs observed in Newer cells. This shift from chemoorganoheterotrophs to chemolithoautotrophs was also observed by Sekhohola-Dlamini *et al.* (2020), who found that a 14-year old MSW landfill with higher total carbon and N was dominated by copiotrophs, while a nearby 36-year old landfill was dominated by lithotrophs (13). Taxonomic shifts were also reflected in differences in functional gene abundances between Older and Newer cells. As noted above, relative abundance of MAGs with genes associated with the fermentation of carbon sources was significantly lower in Older cells. Possibly associated with increases in oxygen availability, significantly more MAGs in Older cells had genes associated with electron transport chain complexes I-IV, indicating the potential for aerobic respiration under changing redox conditions. Furthermore, Older cells had more metabolically flexible microbial communities, with significantly higher relative abundance of MAGs with genes for reduction and/or oxidation of Fe, N, and S. For example, members of the genera *Gallionella* and *Sideroxyarcus* (formerly *Sideroxydans*), which were enriched in Older cells, are widely known to oxidize ferrous iron (36–38). Similarly, members of two other enriched genera, *Sulfuricurvum* and *Sulfurimonas*, contain members known to be metabolically flexible, capable of utilizing a variety of reduced S compounds as electron donors and nitrate or oxygen as electron acceptors (39–41). Concentrations of nitrate, nitrite, and sulfide in leachate were all relatively low (generally < 5 mg/L across all time points), indicating either low levels available in the landfilled waste, steady conversion/uptake rates by the microbial community, or both. In contrast, concentrations of sulfate were relatively higher (> 10 mg/L) throughout the landfill life cycle, with no significant differences between Newer and Older cells at the time of sampling.

### 3.2 Redox response across decades of landfill waste decomposition

Ammonia-N, sulfate, and Fe concentrations in leachate spiked at different points in the landfill life cycle. Each of these redox-associated metabolites was linked to different biogeochemical cycles in our dataset. First, other studies have reported a spike in ammonia-N midway through the landfill life cycle as organics are depleted, mobilizing N via hydrolysis of proteins in OM (3, 5). We observed a similar trend, with ammonia-N broadly peaking with conductivity, bicarbonate, potassium, and sodium at around 20 years of landfill waste age (Figure 5b - cluster 4). This peak in conductivity, bicarbonate, and base cations (K and Na) has been observed elsewhere. Conductivity has been observed to increase in young and intermediate-aged landfills as ions are released from breakdown of organic material (3, 6, 42). Contributing to conductivity, bicarbonate tends to increase in leachate as degradation of organic material produces CO_2_, forming carbonic acid and ultimately, bicarbonate ions (42, 43).

A notable trend during the mobilization and then reduction in labile carbon substrates from 0 – 10 years was an increase in leachate concentrations of Fe, Ca, Cd, and Cr, with peaks occurring around 6-8 years. (Figure 5b – Cluster 2). Although mobilization of these elements may be linked to their pH and redox-dependent natures (3, 8), it is important to note that pH generally remained above 6 in all samples at all time points, a phenomenon observed in other landfills’ anaerobic acid phase (14). It is therefore unlikely that mobilization of Fe, Ca, Cd, and Cr is entirely pH-dependent, as the measured pH was not acidic enough to release these metals into solution. Instead, it is possible that metals formed soluble complexes or colloids with OM, increasing leachate concentrations relative to the later parts of the landfill life cycle (3). Furthermore, reducing conditions in the anaerobic phases may have influenced metal mobility, reducing, for example, Fe(III) to soluble Fe(II) (14). Further study is needed to fully understand the links between organic chemistry and metal cycling in landfill leachate over time.

Finally, concentrations of sulfate, Cu, Mg, Ni, and Zn, were all linked to the historic practice of leachate recirculation at the site (Figure 4; Figure 5b). Leachate recirculation is common in MSW landfills, where it is used to raise moisture content and hasten anaerobic decomposition (3, 44). At our site, leachate recirculation was practiced until the end of 2009, with variable volumes recirculated in each cell monthly. Linear regression analysis revealed that the volume of leachate recirculated was negatively associated with redox and nitrite-N and positively with BOD, COD, sulfate, Cu Cr, Fe, Mg, Ni, and Zn over time. Outside of studies specifically investigating leachate recirculation in landfills (44, 45), few mention if the practice was performed at their sites or consider the broader impacts recirculation has on microbial guilds that control waste decomposition, making it difficult to put our results in context. From our data, we suggest that more reducing conditions (decreases in redox potential) are an expected response to leachate recirculation, which is broadly associated with increased rates of fermentation of organic matter, possibly leading to higher BOD and COD (44, 45). Increased moisture content, brought about by recirculation of leachate, can also promote dissolution of metal-bearing minerals (*e.g.*, Fe/Mn-oxides) and mobilize metals previously bound to solid phases – this is particularly true for metals like Cu, Ni, and Zn that more readily solubilize or bind to organic ligands, thus keeping them in the leachate solution (14). Increased moisture content in landfill cells may also increase contact time, which can release a variety of ions from solid phases, including metals and sulfate.

### 3.2 Metal fate within a landfill

Given concerns about the impact of increasingly oxidizing conditions on metal mobility and metal leaching in landfills (4, 14), one of the goals of our study was to investigate changes in metal availability and putative metal cycling over the landfill life cycle. As mentioned above, the practice of leachate recirculation appeared to accelerate metal release into landfill leachate. Our study identified a previously-unreported association between leachate recirculation and metal concentrations, allowing an initial examination of this phenomenon. Leachate recirculation ceased at this site in 2009, so we had more data for leachate recirculation from Older and Intermediate cells (A, B, C, D1, D2), with cell E only filling from 1999 onwards, and cell F not extant in 2009 (2014 start to filling). Overwhelmingly, metal concentrations were higher in A, B, and C (all filled before 1993) at the time of filling than in D1 and D2 (filled between 1993 and 1998), possibly due to changes in waste streams and waste management practices over time. More research is needed to investigate the impacts of leachate recirculation on metal mobility and fate, as existing research focuses on recirculation’s role in accelerating breakdown of OM.

We hypothesized that Older landfill cells would host microbial communities with a higher abundance of functional genes associated with utilization of inorganic electron acceptors and cycling of metals. We found that the abundance of metal resistance genes was significantly higher in Older cells relative to Newer. Many different genes for metal resistance (for As, Cd, Cu, Co, Cr, Fe, Ni, Se, Te) were significantly more abundant in Older cells than Newer cells, despite lower metal concentrations. Genes for Fe oxidation and reduction were the only metal metabolism genes with significantly higher proportional abundances in Older cells. Interestingly, outside of Fe-cycling, the majority of metal resistance and metal cycling genes or gene groups were encoded by lower abundance MAGs that were not part of the most dominant families in Older cells (Gallionellaceae and Sulfurimonadaceae). Genes for As resistance were some of the most ubiquitous in Older cells, present across a variety of taxa including many, but not all, Gallionellaceae and Sulfurimonadaceae MAGs, lineages which harbour As resistance genes in other environments (40, 46). Genes for As resistance have been found across a variety of environments (21), including deep groundwater systems (46) and permafrost (47). While metal exposure may partially explain the high incidence of metal resistance genes in Older cells, which had highest metal levels historically (48), some studies have seen little overlap between metal exposure and relevant metal-resistance genes (46, 49). It is possible that fluctuations in redox conditions within the solid material in Older landfill cells results in spatial differences in metal exposure that influence microbial community composition and functional gene profiles, but that is difficult to determine without sampling solid landfill waste. Our study provided a comprehensive look at a wide range of metals, and further study is needed to elucidate direct and indirect biogeochemical cycling pathways for landfill metals.

### 3.3 Conclusions

In our study, we examined microbial communities, functional genes, and leachate chemistry across a landfill’s cells of varying ages (2-39 years). We also tracked changes in geochemical conditions across 36 years of landfill leachate chemistry data. We found that the early phases of the landfill life cycle were dominated by fermentative processes, with labile carbon declining over time. Conditions became more oxidizing over time, with evidence of oxygen buildup in Older cells. Microbial diversity declined with time, and there was a clear shift from fermentative processes to utilization of alternative electron donors and acceptors. Our results were largely consistent with studies that modelled potential biogeochemical cycles in landfills in the stabilization phase, having never recorded them (3, 4).

Leachate metal concentrations were influenced by OM content and historic leachate recirculation practices. Leachate recirculation was associated with increases in metal mobility and the potential for metal leaching out of the system, possibly linked to organo-metal complexes. Future research investigating organo-metal complexes and the ability of natural landfill decomposition to pull metals into solution would be an exciting new avenue to better understand metal fate in these environments, especially in conjunction with better modeling for how leachate recirculation-modified moisture regimes impact metal leaching and OM cycling.

Our conceptual model of the key processes and metabolisms over a landfill’s life cycle provides a new framework, based on empirical data, and synthesizing physical processes with microbial impacts. We note that this landfill’s cells were only at the beginning of the stabilization phase. The landfill should continue to be monitored for changes in geochemical conditions and microbial community succession over time to better understand the unexplored dynamics of this crucial, final stage in a landfill life cycle.

## 4.0 Materials and Methods

### 4.1 Site Description

The sampled site is a municipal landfill located in the northeastern United States (NEUS). The site is composed of six landfill cells (A, B, C, D, E, and F), two of which are divided into sub-cells (D into D1 and D2, and F into F1 and F2), each equipped with leachate collection systems. Previous work at this site classified the landfill cells using a five-phase conceptual model based on leachate chemistry and biogas data (1). Cell A, which received waste from 1980 to 1982, and Cell B, which received waste from 1982 to 1988, were both classified as in phase 5. Cell C, which received waste from 1988 – 1993 was classified into phase 4. The authors of this prior study identified evidence of oxygen intrusion into cells A and B, informing their classification into phase 5 of the landfill life cycle. The two subcells of cell D were filled at different times. D1 received waste from 1993 – 1998 and cell D2 from 1995 – 1998. Both were classified into phase 4, due primarily to low gas production relative to cells E and F. Cell E was filled from 1999 – 2014 and was classified into phase 3 (rapid methanogenesis) due to high gas output, mostly composed of methane. Cell F1 filled from 2014 up to the time of sampling (March 2019) and F2 from 2016 up to the time of sampling (March 2019). Both were identified as transitioning from phase 2 (anaerobic acid phase) to phase 3 due to variable gas production but still high proportions of methane (1). For this study, decreases in oxygen demand (BOD) and chemical oxygen demand (COD) at A, B, and to a lesser extent, C, relative to the other cells prompted us to classify A, B, and C as “Older”. Cells E, F1, and F2 were categorized as “Newer”, while D1 and D2 were labelled “Intermediate”.

### 4.2 Leachate sampling

In 2019, we collected leachate samples from each cell/sub-cell for shotgun metagenomic sequencing. One leachate monitoring well at each cell was purged of 2-3 well volumes, and one litre of leachate was collected using a peristaltic pump. Leachate samples were stored on ice and transported to the lab before being filtered through a 0.22 μm Sterivex filter (Millipore Sigma, Burlington, MA) on the day of sampling. Filters were stored at −80°C until DNA extractions were performed. Detailed descriptions of DNA extraction, sequencing, filtering, assembly, binning, and taxonomic assignment can be found in Grégoire *et al.* 2023 (1), which exclusively explored the predicted methane-cycling populations across the landfill. The pipeline for microbial community analyses is described in brief below.

### 4.3 DNA sequencing and prior analyses

DNA was extracted from biomass on Sterivex filters using the PowerSoil DNA Isolation Kit (Qiagen) according to the manufacturer’s protocols but with the filter added to the initial bead tube in place of soil, and with elution into 50 µL of final buffer. DNA was quality-checked using a NanoDrop 1000 (Thermo Scientific, Waltham, MA) and quantified using Qubit fluorimetry (Thermo Scientific, Waltham, MA). Extracted DNA was sent to The Centre for Applied Genomics in Toronto, Canada for shotgun metagenomic sequencing on an Illumina HiSeq machine with paired 2x150 bp reads. Acquired metagenomic reads were quality-trimmed with bbduk in the BBTools suite (https://sourceforge.net/projects/bbmap/) and Sickle v1.33 (50) prior to assembly using SPAdes3 v3.15.5 (51). Scaffolds were trimmed by length (≥ 2.5 kbp) and reads were mapped onto scaffolds using Bowtie2 v2.3.4.1 (52). Scaffolds were binned using three different binning algorithms (CONCOCT v0.4.0, MaxBin2 v2.2.6, MetaBAT2 v2.12.1) and bins were dereplicated using DAS Tool v 1.1.1 before quality-checking in CheckM v 1.0.13 (53–57). A total of 1,881 metagenome-assembled genomes (MAGs) were retained with >70% completion and <10% contamination. Relative abundance of MAGs was calculated using the mean coverage for unique scaffold identifiers within each genome bin. To evaluate the functional potential of microbial communities, MAGs were annotated using Distilled and Refined Annotation of Metabolism (DRAM) v1.0 (19) to look at a wide array of functional genes using both sequence similarity and hidden Markov model (HMM) based detection.

### 4.4 Data analyses

All previous work by Grégoire *et al.* is outlined in brief in section 4.3 (1). For this study, taxonomy was assigned to metagenome-assembled genomes (MAGs) using the Genome Taxonomy Database Toolkit (GTDB-tk) r266 (15). Phylogenomic analysis for all MAGs was performed using GToTree v1.6, using a suite of 25 hidden Markov models (HMMs) for bacteria and archaea to identify single copy genes and generate a tree (58). 1,570 out of our 1,647 MAGs were placed on the tree. The resultant tree was visualized using the Interactive Tree of Life tool (59). To investigate similarity of MAGs across landfill cells, we utilized the dereplication tool dRep, which uses Mash and genome-wide ANI (gANI) to create clusters of MAGs based on pairwise comparisons, choosing a similarity cutoff of 99% (60).

The product file from DRAM was used to investigate the presence of genes associated with central metabolism, carbon-cycling, nitrogen-cycling, and sulfur-cycling. DRAM combines genes into functional gene groups based on the presence of key genes. For example, for the function “acetate, pt 1” to be considered present, MAGs need to have genes for both phosphate acetyltransferase and acetate kinase. We utilized these DRAM categories in all cases except the SOX complex for oxidation of thiosulfate, which DRAM considers to be present if any *sox* gene (*A, B, C, D, X, Y,* or *Z)* is present. Researchers have demonstrated that the presence of *soxXA, soxB,* and *soxYZ* are necessary for partial oxidation of thiosulfate, with *soxCD* completing the reaction (61). Accordingly, we decided to apply a stricter threshold than DRAM, considering the combined presence of *soxA, soxB, soxX, soxY,* and *soxZ* as required to indicate the ability for partial or complete oxidation of thiosulfate.

DRAM also annotates metabolic pathways and multi-subunit complexes (*e.g.*, electron transport chain [ETC]) based on the proportion of genes present. We considered a pathway or complex to be present when over 70% of genes were present. We utilized HMM-based pipelines for Fe and As metabolism and resistance, using FeGenie (20) for iron and Dunivin *et al*.’s arsenic resistance and metabolism pipeline (21). FeGenie uses a library of curated HMMs to group iron-related functional genes, while Dunivin *et al.*’s pipeline looks for individual marker genes using their HMM set. We also utilized BacMet version 2.0 for targeted annotation of metal-resistance genes by looking for homologs to experimentally-confirmed genes from the BacMet database 22). Genes from the BacMet database were filtered to only those which were involved in metal resistance, excluding genes related to genes for antibiotic resistance. Because BacMet and the Dunivin *et al*. pipeline employ different methods, we retained hits from both, even if the genes overlapped (*e.g.*, *arsB* from Dunivin vs. *arsB* from BacMet). Further information on genes contributing to functional groups can be found in the documentation for the tools listed above (Table S2).

All subsequent data analyses, including statistics and generation of figures, were performed in R version 4.3.1 (2023-06-16 ucrt) (62). We separated data into three categories for downstream analysis: Older cells aged 31 – 39 years (cells A, B, and C) and Newer cells aged 3 – 20 years (cells E, F1, and F2). D1 and D2 (aged 24 – 26) were considered Intermediate and were not included in statistical analyses. Base R was used for statistical tests. The packages “phyloseq” and “microeco” were used for generating Venn diagrams of MAG overlap among categories and for producing relative abundance bar graphs (63, 64). Relative abundance was calculated by summing the mean coverage for all MAGs at a given taxonomic level (*e.g.*, at the Family level). Similarly, functional gene abundance was calculated using the relative coverage of MAGs that possessed a given functional gene, group, or pathway. Due to the variety of tools used for annotating functional genes, we collapsed data into binary presence-absence data for each gene, pathway, or gene group. Therefore, if 3 MAGs in a cell had coverages of 1%, 2%, and 3%, and each possessed at least one *arsA* gene, the “abundance” of the functional gene would be 6%. Wilcoxon rank sum tests were used to determine differences in functional gene abundance and chemical parameters between Older and Newer cells. Finally, heatmaps of functional gene abundance were generated using the package “ggplot2” (65).

### 4.5 Leachate chemistry

Thirty-six years of leachate chemistry records were provided by the site owners, with all analyses performed by Brickhouse Environmental consultants. While many parameters were provided by the site owners, we selected the following as relevant to microbial function and showing variance over time; pH, oxidation-reduction potential, electrical conductivity, bicarbonate, BOD, COD, TOC, sulfate, sulfide, ammonia-N, nitrate-N, nitrite-N, organic acids (acetic acid, propionic acid, isobutyric acid, butyric acid, isovaleric acid, valeric acid), total phenolics, base cations (Ca, K, Mg, Na) and metals (As, Cd, Co, Cr, Cu, Fe, Mn, Ni, Se). As leachate chemistry has changed greatly over time for some cells, we reported chemistry both during the time of filling and in the 5 years prior to our sampling to capture shifts over time.

To investigate trends in geochemical parameters over time, we computed 5-year rolling averages across all cells by time since filling across all 36 years (year 0 – 36) of provided leachate data. Then, we computed z-scores for each parameter, transforming data to a mean of 0 and standard deviation of 1 so trends across parameters could be compared agnostic of units.

Using an elbow plot of within-cluster sum of squares to determine the optimum numbers of clusters (Figure S4), we clustered the z-score adjusted data into 8 k-means clusters (Table 2). Plots of 5-year rolling averages were developed using the “ggplot2” package in R (65). Finally, to investigate the impact of the historic practice of leachate recirculation on leachate geochemistry, we ran linear regressions on z-score scaled 5-year rolling averages for all cells across the time window for which there was active leachate recirculation (> 5 gallons of leachate/month).

## Acknowledgements

We are sincerely grateful to the landfill site management (anonymity by request) and Brickhouse Environmental consulting for site access, aid with sampling, and provision of detailed monitoring records. We thank Dr. Jennifer Biddle and her lab group for hosting our team for sample processing prior to shipment. Thanks also to Dr. Nikhil George, Ms. Rebecca Co, and Dr. Alexandra Sauk for help with sampling, and to Dr. Nikhil George for his work with binning pipelines. This work was supported by an NSERC Discovery Grant (2016–03686) and an Ontario Early Research Award (ER19-15-228) to LAH. LAH was supported by a Tier II Canada Research Chair.

## Data Availability

All data involved in this research are publicly available. Sequencing data have been deposited to NCBI under the BioProject <u>PRJNA900590</u>, with raw reads available on the SRA database. Geochemical data, microbial functional annotations, and microbial taxonomy data are all available in the Open Science Framework (https://doi.org/10.17605/OSF.IO/MSDB8). All R code used for statistical analyses and to generate figures are also on the Open Science Framework.

## CRediT authorship contribution statement

**Kimber E. Munford:** Data curation, formal analysis, investigation, methodology, visualization, writing – original draft, writing – review & editing. **Daniel S. Grégoire:** Data curation, formal analysis, investigation, methodology, software, writing – review & editing. **Laura A. Hug:** Conceptualization, data curation, funding acquisition, project administration, resources, software, supervision, writing – review & editing.

