## Supplementary materials for "Tracking interlinked microbial and geochemical succession over decades in landfilled municipal solid waste"

**Table S1: Relative abundance of MAGs within the most abundant families in Older and Newer samples in an aging landfill.** Taxonomy is taken from GTDB-tk r266, where lineages starting UBA or consisting of a string of letters and numbers (e.g. JAQTOM01) are officially recognized but not yet formally named. dRep results that are the same (e.g. 214_1 and 214_1) represent 99% similarity.

| **Family** | **Genus** | **Dereplicated Group** | **MAG number** | **Cell** | **Relative Abundance (%)** |
| --- | --- | --- | --- | --- | --- |
| Gallionellaceae | *Gallionella* | 270_0 | 17 | A | 11.7 |
|  | *Gallionella* | 214_2 | 49 | B | 10.8 |
|  | *Gallionella* | 214_1 | 38 | C | 1.91 |
|  | *Gallionella* | 214_1 | 53 | B | 0.613 |
|  | *Gallionella* | 268_0 | 67 | B | 9.21 |
|  | *Gallionella* | 267_1 | 7 | C | 8.57 |
|  | *Gallionella* | 267_1 | 61 | B | 0.329 |
|  | *Gallionella* | 267_1 | 186 | D1 | 0.142 |
|  | *Gallionella* | 271_0 | 52 | A | 8.08 |
|  | *Gallionella* | 269_0 | 51 | B | 2.70 |
|  | *Gallionella* | 273_0 | 75 | F1 | 0.371 |
|  | *Gallionella* | 262_0 | 80 | C | 0.257 |
|  | *Gallionella* | 272_0 | 117 | C | 0.242 |
|  | *Sideroxyarcus* | 265_0 | 48 | B | 1.81 |
|  | *Sideroxyarcus* | 261_0 | 72 | B | 1.16 |
|  | *Sideroxyarcus* | 263_1 | 108 | C | 0.394 |
|  | *Sideroxyarcus* | 263_1 | 61 | D1 | 0.185 |
|  | *Sideroxyarcus* | 264_0 | 76 | C | 0.141 |
|  | JAQTOM01 | 266_0 | 88 | A | 4.62 |
| Sulfurimonadaceae | *Sulfuricurvum* | 1105_1 | 16 | A | 15.8 |
|  | *Sulfuricurvum* | 1105_1 | 99 | B | 0.247 |
|  | *Sulfuricurvum* | 1102_0 | 65 | A | 3.35 |
|  | *Sulfuricurvum* | 1103_0 | 35 | B | 1.52 |
|  | *Sulfuricurvum* | 1104_1 | 6 | B | 0.922 |
|  | *Sulfuricurvum* | 1104_1 | 55 | C | 0.131 |
|  | *Sulfuricurvum* | 1096_0 | 37 | C | 0.903 |
|  | *Sulfuricurvum* | 1098_0 | 110 | E | 0.774 |
|  | *Sulfuricurvum* | 1101_1 | 144 | B | 0.346 |
|  | *Sulfuricurvum* | 1101_1 | 10 | C | 0.0950 |
|  | *Sulfuricurvum* | 1099_1 | 25 | E | 0.338 |
|  | *Sulfuricurvum* | 1099_1 | 21 | C | 0.116 |
|  | *Sulfuricurvum* | 1099_2 | 85 | B | 0.162 |
|  | *Sulfuricurvum* | 1097_0 | 133 | D1 | 0.335 |
|  | *Sulfuricurvum* | 1106_0 | 91 | A | 0.180 |
|  | *Sulfuricurvum* | 1100_1 | 34 | C | 0.134 |
|  | *Sulfurimonas* | 1109_0 | 8 | A | 5.84 |
|  | *Sulfurimonas* | 1117_0 | 9 | E | 1.67 |
|  | *Sulfurimonas* | 1110_1 | 138 | E | 1.58 |
|  | *Sulfurimonas* | 1112_0 | 89 | E | 0.438 |
|  | *Sulfurimonas* | 1113_2 | 28 | A | 0.409 |
|  | *Sulfurimonas* | 1113_1 | 12 | A | 0.347 |
|  | *Sulfurimonas* | 1107_1 | 122 | C | 0.334 |
|  | *Sulfurimonas* | 1107_1 | 101 | B | 0.159 |
|  | *Sulfurimonas* | 1107_1 | 56 | D2 | 0.0747 |
|  | *Sulfurimonas* | 1116_0 | 10 | E | 0.241 |
|  | *Sulfurimonas* | 1108_0 | 53 | C | 0.193 |
|  | *Sulfurimonas* | 1111_0 | 76 | E | 0.174 |
|  | *Sulfurimonas* | 1114_0 | 9 | F1 | 0.115 |
|  | *Sulfurimonas* | 1115_0 | 61 | E | 0.106 |
| Arcobacteraceae | *Aliarcobacter* | 1122_1 | 123 | C | 27.2 |
|  | *Aliarcobacter* | 1122_2 | 124 | C | 16.7 |
|  | *Aliarcobacter* | 1120_2 | 67 | E | 5.19 |
|  | *Aliarcobacter* | 1120_3 | 88 | D2 | 1.35 |
|  | *Aliarcobacter* | 1120_1 | 71 | D2 | 0.523 |
|  | *Aliarcobacter* | 1121_1 | 58 | D2 | 3.35 |
|  | *Aliarcobacter* | 1121_2 | 119 | F1 | 0.548 |
|  | *Aliarcobacter* | 1123_0 | 80 | D2 | 0.990 |
|  | CAIJNA01 | 1118_0 | 79 | C | 0.495 |
| Dysgonomonadaceae | *Proteiniphilum* | 547_1 | 34 | F1 | 5.33 |
|  | *Proteiniphilum* | 547_1 | 43 | F2 | 3.32 |
|  | *Proteiniphilum* | 547_1 | 40 | E | 2.01 |
|  | *Proteiniphilum* | 547_1 | 192 | D2 | 0.359 |
|  | *Proteiniphilum* | 547_1 | 65 | C | 0.291 |
|  | *Proteiniphilum* | 547_1 | 196 | D1 | 0.246 |
|  | *Proteiniphilum* | 547_1 | 37 | A | 0.203 |
|  | *Proteiniphilum* | 536_1 | 159 | F2 | 5.15 |
|  | *Proteiniphilum* | 536_1 | 145 | F1 | 0.418 |
|  | *Proteiniphilum* | 536_1 | 237 | D2 | 0.160 |
|  | *Proteiniphilum* | 536_1 | 60 | E | 0.142 |
|  | *Proteiniphilum* | 536_1 | 109 | C | 0.139 |
|  | *Proteiniphilum* | 544_1 | 60 | F2 | 5.11 |
|  | *Proteiniphilum* | 544_1 | 49 | D2 | 1.80 |
|  | *Proteiniphilum* | 544_1 | 45 | F1 | 1.22 |
|  | *Proteiniphilum* | 544_1 | 105 | C | 0.134 |
|  | *Proteiniphilum* | 544_1 | 12 | E | 0.114 |
|  | *Proteiniphilum* | 537_1 | 15 | F2 | 1.34 |
|  | *Proteiniphilum* | 537_1 | 279 | F1 | 0.0794 |
|  | *Proteiniphilum* | 534_1 | 77 | F2 | 0.980 |
|  | *Proteiniphilum* | 534_1 | 67 | F1 | 0.112 |
|  | *Proteiniphilum* | 546_1 | 107 | F2 | 0.524 |
|  | *Proteiniphilum* | 546_1 | 201 | F1 | 0.157 |
|  | *Proteiniphilum* | 535_1 | 201 | E | 0.488 |
|  | *Proteiniphilum* | 535_1 | 124 | F2 | 0.328 |
|  | *Proteiniphilum* | 548_1 | 196 | E | 0.284 |
|  | *Proteiniphilum* | 548_1 | 58 | F2 | 0.213 |
|  | *Proteiniphilum* | 545_0 | 185 | F2 | 0.149 |
|  | *Proteiniphilum* | 538_0 | 118 | F2 | 0.126 |
|  | *Proteiniphilum* | 548_1 | 4 | F1 | 0.123 |
|  | *Petrimonas* | 539_1 | 226 | F2 | 2.00 |
|  | *Petrimonas* | 539_1 | 47 | E | 0.472 |
|  | *Petrimonas* | 539_1 | 276 | F1 | 0.392 |
|  | *Petrimonas* | 539_1 | 104 | D2 | 0.104 |
|  | *Petrimonas* | 539_1 | 118 | C | 0.0811 |
|  | *Petrimonas* | 540_0 | 36 | E | 1.65 |
|  | UBA4179 | 541_1 | 69 | E | 3.20 |
|  | UBA4179 | 541_1 | 93 | F2 | 0.0914 |
|  | UBA5287 | 549_1 | 83 | F2 | 1.25 |
|  | UBA5287 | 549_1 | 194 | D2 | 0.159 |
|  | UBA5287 | 549_1 | 69 | F1 | 0.123 |
|  | UBA2632 | 533_1 | 176 | E | 0.348 |
|  | UBA2632 | 533_1 | 155 | D2 | 0.234 |
|  | UBA2632 | 533_1 | 172 | F1 | 0.196 |
|  | UBA2632 | 533_1 | 206 | F2 | 0.176 |
|  | UBA2632 | 1002_0 | 177 | F2 | 0.0842 |
|  | JAQVEB01 | 542_0 | 125 | E | 0.105 |
| Cloacimonadaceae | UBA5456 | 510_1 | 112 | F2 | 9.42 |
|  | UBA5456 | 510_1 | 175 | F1 | 7.79 |
|  | UBA5456 | 510_1 | 117 | D2 | 1.80 |
|  | UBA5456 | 510_1 | 43 | E | 0.816 |
|  | UBA5456 | 510_1 | 67 | C | 0.771 |
|  | UBA5456 | 510_1 | 30 | D1 | 0.201 |
|  | UBA5456 | 510_1 | 42 | A | 0.194 |
|  | UBA5456 | 512_1 | 267 | F1 | 4.17 |
|  | UBA5456 | 512_1 | 150 | F2 | 1.85 |
|  | UBA5456 | 512_1 | 4 | D2 | 0.649 |
|  | UBA5456 | 512_1 | 84 | C | 0.185 |
|  | UBA5456 | 512_1 | 26 | E | 0.154 |
|  | UBA5456 | 513_1 | 105 | F2 | 0.720 |
|  | UBA5456 | 513_1 | 237 | F1 | 0.301 |
|  | UBA5456 | 511_1 | 216 | F2 | 0.211 |
|  | UBA5456 | 511_1 | 209 | F1 | 0.0524 |
|  | UBA5456 | 509_1 | 148 | F2 | 0.202 |
|  | UBA5456 | 509_1 | 173 | F1 | 0.139 |
|  | UBA5456 | 509_1 | 21 | D2 | 0.0820 |
|  | UBA5456 | 514_0 | 55 | E | 0.196 |
|  | UBA5456 | 520_0 | 5 | E | 0.144 |
|  | UBA5456 | 518_1 | 51 | E | 0.121 |
|  | UBA5456 | 518_1 | 29 | D2 | 0.0902 |
|  | UBA5456 | 519_2 | 146 | F1 | 0.114 |
|  | UBA5456 | 519_1 | 35 | E | 0.112 |
|  | *Syntrophosphaera* | 503_1 | 190 | E | 1.75 |
|  | *Syntrophosphaera* | 503_1 | 197 | D1 | 1.29 |
|  | *Syntrophosphaera* | 508_0 | 8 | E | 1.08 |
|  | *Syntrophosphaera* | 507_0 | 180 | F1 | 0.151 |
|  | *Syntrophosphaera* | 506_0 | 31 | D2 | 0.100 |
|  | *Cloacimonas* | 505_1 | 95 | D1 | 0.718 |
|  | *Cloacimonas* | 505_1 | 21 | E | 0.640 |
|  | *Cloacimonas* | 504_1 | 144 | E | 0.369 |
|  | *Cloacimonas* | 504_1 | 144 | F2 | 0.266 |
|  | *Cloacimonas* | 504_1 | 87 | F1 | 0.0574 |
|  | *Cloacimonas* | 516_0 | 18 | D2 | 0.0844 |
|  | *Cloacimonas* | 517_0 | 13 | F1 | 0.0565 |

**Table S2: Wilcoxon rank sum test results for relative abundance of all MAGs containing functional genes, gene groups, or pathways of interest.** Only results which were significant at p < 0.05 are reported. Bolded values are those which were no longer significantly different after the Benjamini-Hochberg (BH) correction.

| **Category** | **Element** | **Source** | **Functional group, gene, or pathway** | **W** | **p-value** | **BH corrected p-value** |
| --- | --- | --- | --- | --- | --- | --- |
| Central C metabolism | C | DRAM | Reductive pentose phosphate cycle^1^ | 134921 | < 0.001 | < 0.001 |
|  | C | DRAM | Pentose phosphate cycle^1^ | 134847 | < 0.001 | < 0.001 |
|  | C | DRAM | Reductive citrate cycle^1^ | 132542 | < 0.001 | < 0.001 |
|  | C | DRAM | Glyoxylate cycle^1^ | 143780 | < 0.001 | < 0.001 |
|  | C | DRAM | Citrate cycle^1^ | 120627 | < 0.001 | < 0.001 |
|  | C | DRAM | Entner-Doudoroff pathway^1^ | 147554 | 0.0269 | **0.0603** |
| Carbon cycling: CAZy | C | DRAM | Alpha-mannan (backbone and oligo cleavage)^2^ | 159563 | 0.0133 | 0.0334 |
|  | C | DRAM | Arabinan (backbone and oligo cleavage)^2^ | 169819 | < 0.001 | < 0.001 |
|  | C | DRAM | Arabinose cleavage (oligo cleavage) ^2^ | 187613.5 | < 0.001 | < 0.001 |
|  | C | DRAM | Beta-galactan (pectic galactan) (backbone and oligo cleavage) ^2^ | 159232 | 0.0119 | 0.0304 |
|  | C | DRAM | Beta-mannan (backbone and oligo cleavage) ^2^ | 163259.5 | 0.0219 | **0.0527** |
|  | C | DRAM | Fucose Cleavage (oligo cleavage) ^2^ | 178634 | < 0.001 | < 0.001 |
|  | C | DRAM | Mixed-Linkage glucans (backbone and oligo cleavage) ^2^ | 140782 | 0.00429 | 0.0122 |
|  | C | DRAM | Pectin (backbone and oligo cleavage) ^2^ | 170935.5 | < 0.001 | < 0.001 |
|  | C | DRAM | Polyphenolics (oxidative enzymes: laccases and peroxidases)^2^ | 134886.5 | < 0.001 | < 0.001 |
|  | C | DRAM | Rhamnose cleavage (oligo cleavage) ^2^ | 177604 | < 0.001 | < 0.001 |
|  | C | DRAM | Starch (backbone and oligo cleavage) ^2^ | 166331 | 0.00530 | 0.0144 |
|  | C | DRAM | Sulf-Polysachharides (backbone and oligo cleavage) ^2^ | 165849 | < 0.001 | < 0.001 |
|  | C | DRAM | Xyloglucan (backbone and oligo cleavage) ^2^ | 168752 | < 0.001 | 0.00137 |
| Carbon cycling: SCFA and alcohol conversions | C | DRAM | Butyrate, pt 1 (phosphate butyryltransferase) | 166950 | < 0.001 | < 0.001 |
|  | C | DRAM | Butyrate, pt 2 (butyrate kinase and medium-chain acyl-CoA synthetase) | 193226 | < 0.001 | < 0.001 |
|  | C | DRAM | acetate, pt 1 (phosphate acetyltransferase and acetate kinase) | 171566.5 | < 0.001 | < 0.001 |
|  | C | DRAM | acetate, pt 3 (acetyl-CoA hydrolase) | 152190 | 0.0234 | **0.0551** |
|  | C | DRAM | pyruvate => acetyl CoA v1 (2-oxoglutarate/2-oxoacid ferredoxin oxidoreductase subunit alpha) | 131301 | < 0.001 | < 0.001 |
|  | C | DRAM | pyruvate => acetylCoA + formate v3 (formate C-acetyltransferase) | 176851 | < 0.001 | < 0.001 |
| Carbon cycling: Methane | C | DRAM | *mcrA* gene | 160287 | 0.00132 | 0.00426 |
|  | C | DRAM | acetate => methane, pt 1 (acetyl-CoA synthetase) | 113649 | < 0.001 | < 0.001 |
|  | C | DRAM | acetate => methane, pt 3 (phosphate acetyltransferase) | 171566.5 | < 0.001 | < 0.001 |
|  | C | DRAM | methanol => methane (*mtaB* gene) | 156502 | 0.0242 | **0.0557** |
|  | C | DRAM | putative but not defining CO2 => methane (formylmethanofuran dehydrogenase) | 148900.5 | 0.0244 | **0.0557** |
|  | C | DRAM | methane => methanol, with oxygen (mmo) (*mmoX* gene) | 148600 | < 0.001 | < 0.001 |
|  | C | DRAM | methane => methanol, with oxygen (pmo) (*pmoA* gene) | 147166.5 | < 0.001 | < 0.001 |
| Electron transport chain |  | DRAM | Complex I: NADH:quinone oxidoreductase | 118801 | < 0.001 | < 0.001 |
|  |  | DRAM | Complex II: Succinate dehydrogenase | 140781.5 | < 0.001 | 0.00119 |
|  |  | DRAM | Complex III: Cytochrome bc1 complex respiratory unit | 122708.5 | < 0.001 | < 0.001 |
|  |  | DRAM | Complex III: Cytochrome bd ubiquinol oxidase | 141042 | < 0.001 | < 0.001 |
|  |  | DRAM | Complex IV High affinity: Cytochrome bd ubiquinol oxidase | 141042 | < 0.001 | < 0.001 |
|  |  | DRAM | Complex IV High affinity: Cytochrome c oxidase, cbb3-type | 138307 | < 0.001 | < 0.001 |
|  |  | DRAM | Complex IV Low affinity: Cytochrome c oxidase, prokaryotes | 143008 | < 0.001 | < 0.001 |
|  |  | DRAM | Complex V: F-type ATPase | 119180.5 | < 0.001 | < 0.001 |
|  |  | DRAM | Complex V: V/A-type ATPase | 172414 | < 0.001 | < 0.001 |
| Redox cycling | N | DRAM | Bacterial/archaeal ammonia oxidation (*amoA* gene) | 147166.5 | < 0.001 | < 0.001 |
|  | N | DRAM | Dissimilatory nitrite reduction to ammonia (*nrfA* gene) | 172302.5 | < 0.001 | < 0.001 |
|  | N | DRAM | nitrate => nitrite (*napA* or *narG* gene) | 134520 | < 0.001 | < 0.001 |
|  | N | DRAM | nitric oxide => nitrous oxide (nitric oxide reductase) | 141184 | < 0.001 | < 0.001 |
|  | N | DRAM | nitrite => nitrate (*nxrA* and *nxrB* genes) | 139219.5 | < 0.001 | < 0.001 |
|  | N | DRAM | nitrite => nitric oxide (respiratory nitrite reductase) | 130998 | < 0.001 | < 0.001 |
|  | N | DRAM | nitrogen => ammonia (*nifH* gene) | 130898 | < 0.001 | < 0.001 |
|  | N | DRAM | nitrous oxide => nitrogen (nitrous-oxide reductase) | 148257 | 0.00276 | 0.00800 |
|  | S | DRAM | thiosulfate => sulfate (*soxA, soxB, soxX, soxY*, and *soxZ* genes) | 148284 | < 0.001 | 0.00114 |
|  | S | DRAM | sulfate => sulfide (*dsrA* or *dsrB* gene) | 145486.5 | < 0.001 | < 0.001 |
|  | S | DRAM | thiosulfate => sulfite (*phsA/psrA* gene) | 150502 | 0.0492 | **0.101** |
|  | Fe | FeGenie | iron oxidation (c*yc1, cyc2, foxABC, foxEYZ, pioABC,* Sulfocyanin) | 128193.5 | < 0.001 | < 0.001 |
|  | Fe | FeGenie | possible iron oxidation and possible iron reduction (m*toAB, cyc2 [cluster 3])* | 150289 | < 0.001 | < 0.001 |
|  | Fe | FeGenie | probable iron reduction (m*trCB, mtrAB, mtoAB-mtrC*) | 152731 | 0.0492 | **0.101** |
| Metal cycling and resistance | As | Dunivin | *arsM* | 145754 | 0.00263 | 0.00800 |
|  | As | Dunivin | *acr3* | 135934.5 | < 0.001 | < 0.001 |
|  | As | Dunivin | *arsC* (thioredoxin-dependent) | 147135 | 0.0109 | 0.0283 |
|  | As | Dunivin | *arsC* (glutaredoxin-dependent) | 142312 | < 0.001 | < 0.001 |
|  | As | Dunivin | *arsD* | 148509 | 0.00154 | 0.00487 |
|  | As | Dunivin | *arsB* | 148612.5 | 0.00452 | 0.0127 |
|  | As | BacMet | *arsB* (BAC0573) | 143742 | < 0.001 | **<** 0.001 |
|  | As | BacMet | *arsH* (BAC0034) | 152191 | 0.0235 | **0.0550** |
|  | As | BacMet | *arsH* (BAC0595) | 152893 | 0.0391 | **0.0842** |
|  | As | BacMet | *pstB* (BAC0316) | 141402 | < 0.001 | < 0.001 |
|  | Cr | BacMet | *chrA* (BAC0026) | 152315 | 0.00273 | 0.00800 |
|  | Cr | BacMet | *chrF* (BAC0029) | 150457.5 | < 0.001 | < 0.001 |
|  | Cr | BacMet | *ruvB* (BAC0355) | 138117 | < 0.001 | < 0.001 |
|  | Co, Cd, Ni | BacMet | *dmeF* (BAC0132) | 152315 | 0.00273 | 0.00800 |
|  | Co, Cd, Ni | BacMet | *czcA* (BAC0119) | 153058 | 0.0204 | 0.0499 |
|  | Cu | BacMet | *copR* (BAC0719) | 151076 | 0.00188 | 0.00584 |
|  | Cu | BacMet | *cop-unnamed* (BAC0750) | 152892 | 0.0389 | **0.0842** |
|  | Hg | BacMet | *merT* (BAC0690) | 152893 | 0.0391 | **0.0842** |
|  | Pb | BacMet | *pbrA* (BAC0298) | 152728 | 0.0487 | **0.101** |
|  | Te | BacMet | *terD* (BAC0389) | 153058 | 0.0204 | 0.0499 |

^1^ Lists of genes contributing to DRAM pathways and multi-subunit-complexes can be found in the DRAM documentation and the DRAM paper (1).

^2^ Multiple genes are accounted for when DRAM annotates carbohydrate-active enzymes. More information can be found in the DRAM documentation and DRAM paper (1).

**Table S3: All geochemical measurements for key variables, collated by landfill cell age class during filling of the cells and during the 5 years prior to sampling in winter 2019.** Wilcoxon rank sum test results and Benjamini-Hochberg corrected p-values are presented for comparisons between only Older and Newer cells in the 5 years prior to sampling. See Table 1 for a subset of only the significantly different variables between Older (31-39 years) and Newer (3-20 years) landfill cells in the 5 years before sampling.

|  | **Older (A, B, C)** | | **Intermediate (D1, D2)** | | **Newer (E, F1, F2)** | **Statistics** | |
| --- | --- | --- | --- | --- | --- | --- | --- |
| **Analyte** | **During filling** | **5 years before sampling** | **During filling** | **5 years before sampling** | **5 years before sampling** | **Wilcoxon rank sum result** | **BH-corrected p - value** |
| pH | 6.57 ± 0.544 | 7.39 ± 0.404 | 6.43 ± 0.488 | 7.15 ± 0.335 | 7.42 ± 0.422 | 0.581 | 0.605 |
| Conductivity (μS/cm) | 7522 ± 3764 | 4665 ± 2083 | 5435 ± 2810 | 7791 ± 2580 | 7789 ± 3629 | < 0.001 | < 0.001 |
| Redox (mV) | ND | -36.1 ± 71.8 | -63.3 ± 66.5 | -67.4 ± 59.4 | -88.8 ± 69.5 | < 0.001 | < 0.001 |
| Bicarbonate (mg/L) | 3914 ± 558 | 1557 ± 774 | 1948 ± 1140 | 2722 ± 979 | 1842 ± 1242 | 0.00209 | 0.00257 |
| BOD_5_ (mg/L) | 9629 ± 9940 | 71.6 ± 194.2 | 3768 ± 2421 | 118 ± 114 | 251 ± 308 | < 0.001 | < 0.001 |
| COD (mg/L) | 14285 ± 13298 | 446 ± 387 | 5609 ± 3536 | 860 ± 379 | 1052 ± 875 | < 0.001 | < 0.001 |
| TOC (mg/L) | 3690 ± 3118 | 115 ± 56.2 | 1710 ± 977 | 252 ± 116 | 272 ± 227 | < 0.001 | < 0.001 |
| Sulfate (mg/L) | 173 ± 223 | 16.3 ± 21.7 | 38.7 ± 45.3 | 11.0 ± 7.45 | 48.1 ± 81.6 | 0.235 | 0.306 |
| Sulfide (mg/L) | NA | 0.845 ± 0.471 | NA | 1.22 ± 0.753 | 5.17 ± 11.4 | < 0.001 | < 0.001 |
| Ammonia-N (mg/L) | 289 ± 222 | 219 ± 123 | 153 ± 129 | 405 ± 167 | 458 ± 306 | < 0.001 | < 0.001 |
| Nitrate-N (mg/L) | 1.58 ± 5.99 | 0.295 ± 0.362 | 0.798 ± 0.722 | 0.239 ± 0.0735 | 0.237 ± 0.0674 | 0.258 | 0.306 |
| Nitrite-N (mg/L) | 1.05 ± 1.18 | 1.81 ± 3.38 | 0.423 ± 0.193 | 5.11 ± 8.40 | 2.44 ± 5.10 | 0.0562 | 0.0916 |
| Acetic acid (mg/L) | 2434 ± 1758 | 12.2 ± 25.1 | 1330 ± 834 | 55.6 ± 62.2 | 61.3 ± 71.4 | < 0.001 | < 0.001 |
| Butyric acid (mg/L) | 1366 ± 1527 | 6.70 ± 10.5 | 533 ± 383 | 12.5 ± 20.0 | 13.5 ± 17.6 | 0.0805 | 0.113 |
| Isobutyric acid (mg/L) | 170 ± 139 | 3.65 ± 4.47 | 92.7 ± 54.0 | 3.59 ± 3.86 | 7.61 ± 12.3 | 0.0632 | 0.0920 |
| Isovaleric acid (mg/L) | 158 ± 107 | 2.51 ± 1.54 | 67.9 ± 26.9 | 2.56 ± 1.38 | 4.98 ± 5.76 | 0.0428 | 0.0741 |
| Propionic acid (mg/L) | 903 ± 618 | 9.99 ± 21.5 | 603 ± 355 | 15.3 ± 23.6 | 34.2 ± 72.4 | 0.00812 | 0.0151 |
| Valeric acid (mg/L) | 518 ± 430 | 2.82 ± 1.68 | 223 ± 149 | 2.76 ± 1.26 | 6.08 ± 7.77 | 0.307 | 0.347 |
| Total phenolics (mg/L) | 3.39 ± 2.40 | 0.173 ± 0.183 | 1.68 ± 1.07 | 0.205 ± 0.179 | 0.835 ± 1.59 | < 0.001 | < 0.001 |
| Arsenic (μg/L) | 34.4 ± 37.1 | 16.9 ± 14.5 | 19.0 ± 6.49 | 28.8 ± 16.2 | 61.4 ± 61.4 | < 0.001 | < 0.001 |
| Cadmium (μg/L) | 18.5 ± 21.9 | 1.03 ± 0.922 | 3.74 ± 5.34 | 1.09 ± 1.07 | 1.38 ± 1.41 | 0.485 | 0.525 |
| Calcium (mg/L) | 550 ± 271 | 67.6 ± 17.3 | 354 ± 183 | 96.4 ± 23.2 | 141 ± 89.3 | < 0.001 | < 0.001 |
| Chromium (μg/L) | 160 ± 410 | 6.17 ± 4.12 | 41.0 ± 29.2 | 24.7 ± 16.4 | 56.8 ± 53.8 | < 0.001 | < 0.001 |
| Cobalt (μg/L) | 35.7 ± 18.1 | 10.2 ± 5.96 | 33.7 ± 48.8 | 15.7 ± 4.40 | 16.6 ± 12.1 | 0.0637 | 0.0920 |
| Copper (μg/L) | 34.9 ± 31.7 | 6.82 ± 10.3 | 10.9 ± 2.88 | 7.31 ± 12.6 | 10.7 ± 17.8 | 0.910 | 0.928 |
| Iron (mg/L) | 849 ± 2960 | 29.4 ± 101 | 416 ± 279 | 36.3 ± 12.1 | 51.8 ± 52.1 | < 0.001 | < 0.001 |
| Manganese (mg/L) | 16.8 ± 11.8 | 0.418 ± 0.494 | 13.3 ± 13.7 | 0.382 ± 0.101 | 1.29 ± 1.51 | < 0.001 | < 0.001 |
| Magnesium (mg/L) | 147 ± 127 | 48.0 ± 23.4 | 96.2 ± 43.3 | 70.1 ± 32.1 | 91.8 ± 42.8 | < 0.001 | < 0.001 |
| Nickel (μg/L) | 212 ± 205 | 38.5 ± 26.5 | 73.5 ± 40.5 | 75.8 ± 39.9 | 110 ± 79.8 | < 0.001 | < 0.001 |
| Potassium (mg/L) | 338 ± 145 | 195 ± 94.4 | 179 ± 143 | 289 ± 101 | 242 ± 137 | 0.00101 | 0.00194 |
| Selenium (μg/L) | 25.4 ± 35.1 | 19.7 ± 33.5 | 13.0 ± 6.49 | 15.1 ± 16.1 | 20.9 ± 22.3 | 0.244 | 0.306 |
| Sodium (mg/L) | 702 ± 421 | 469 ± 226 | 452 ± 326 | 879 ± 330 | 736 ± 422 | < 0.001 | < 0.001 |
| Zinc (mg/L) | 21.3 ± 37.5 | 0.0403 ± 0.236 | 0.544 ± 0.807 | 0.0491 ± 0.0560 | 0.0697 ± 0.0730 | < 0.001 | < 0.001 |

* Cell A was filled from 1980-1982, a period without leachate chemistry data, so data for 1983-1984 were used. Cell B filled from 1982-88 (1983-88 data used), cell C from 1988-93, D1 from 1993-1998, and D2 from 1995-1998.

**Table S4: Linear regressions of leachate geochemical parameters as a function of the volume of leachate recirculated in 5-year rolling averages of 36 years of landfill leachate data.**

| **Parameter** | **Estimated Coefficient** | **Standard Error of the Coefficient** | **t value (estimated coefficient/std. error)** | **r^2^** | **p value** |
| --- | --- | --- | --- | --- | --- |
| **pH (as [OH^-^])** | -1.31 x 10^-12^ | 8.27 x 10^-13^ | -1.59 | 0.134 | 0.132 |
| **Conductivity** | 0.0135 | 0.0190 | 0.709 | 0.0305 | 0.488 |
| **Redox** | -5.04 x 10^-4^ | 6.45 x 10^-5^ | -7.81 | 0.792 | **< 0.001** |
| **Bicarbonate** | 0.00641 | 0.00757 | 0.846 | 0.429 | 0.410 |
| **BOD_5_** | 0.0379 | 0.0172 | 2.21 | 0.234 | **0.0419** |
| **COD** | 0.0498 | 0.0216 | 2.30 | 0.249 | **0.0352** |
| **TOC** | 0.0121 | 0.00573 | 2.11 | 0.218 | 0.0510 |
| **Sulfate** | 5.51 x 10^-4^ | 8.40 x 10^-5^ | 6.55 | 0.729 | **< 0.001** |
| **Sulfide** | -9.30 x 10^-6^ | 1.06 x 10^-5^ | -0.877 | 0.0459 | 0.394 |
| **Ammonia-N** | 7.49 x 10^-4^ | 1.43 x 10^-3^ | 0.523 | 0.0168 | 0.608 |
| **Nitrate-N** | 1.29 x 10^-6^ | 1.19 x 10^-6^ | 1.09 | 0.0693 | 0.291 |
| **Nitrite-N** | -7.89 x 10^-6^ | 3.56 x 10^-6^ | -2.22 | 0.235 | **0.0414** |
| **Acetic acid** | 0.00696 | 0.00560 | 1.24 | 0.0882 | 0.232 |
| **Butyric acid** | 0.00544 | 0.00262 | 2.08 | 0.212 | 0.0543 |
| **Isobutyric acid** | 3.72 x 10^-4^ | 3.45 x 10^-4^ | 1.08 | 0.0676 | 0.298 |
| **Isovaleric acid** | 3.64 x 10^-4^ | 2.94 x 10^-4^ | 1.24 | 0.0878 | 0.233 |
| **Propionic acid** | 0.00288 | 0.00236 | 1.22 | 0.0853 | 0.240 |
| **Valeric acid** | 0.00161 | 0.00103 | 1.56 | 0.132 | 0.138 |
| **Total phenolics** | 1.36 x 10^-6^ | 6.34 x 10^-6^ | 0.214 | 0.00287 | 0.833 |
| **Arsenic** | -1.80 x 10^-7^ | 8.80 x 10^-8^ | -1.34 | 0.101 | 0.199 |
| **Cadmium** | 4.15 x 10^-8^ | 2.68 x 10^-8^ | 1.55 | 0.130 | 0.141 |
| **Calcium** | 1.42 x 10^-3^ | 1.02 x 10^-3^ | 1.40 | 0.109 | 0.182 |
| **Chromium** | 1.71 x 10^-6^ | 6.29 x 10^-7^ | 2.72 | 0.316 | **0.0153** |
| **Cobalt** | -1.16 x 10^-7^ | 8.31 x 10^-8^ | -1.39 | 0.108 | 0.184 |
| **Copper** | 1.19 x 10^-7^ | 4.44 x 10^-8^ | 2.69 | 0.311 | **0.0162** |
| **Iron** | 0.00677 | 0.00141 | 4.80 | 0.590 | **< 0.001** |
| **Manganese** | -2.51 x 10^-5^ | 4.16 x 10^-5^ | -0.604 | 0.0223 | 0.554 |
| **Magnesium** | 5.79 x 10^-4^ | 8.07 x 10^-5^ | 7.17 | 0.763 | **< 0.001** |
| **Nickel** | 1.36 x 10^-6^ | 2.72 x 10^-7^ | 5.01 | 0.610 | **< 0.001** |
| **Potassium** | 4.67 x 10^-4^ | 0.00130 | 0.359 | 0.00800 | 0.724 |
| **Selenium** | -5.44 x 10^-9^ | 1.68 x 10^-8^ | -0.323 | 0.00649 | 0.751 |
| **Sodium** | 0.00237 | 0.00293 | 0.808 | 0.0392 | 0.431 |
| **Zinc** | 1.38 x 10^-4^ | 1.12 x 10^-5^ | 12.3 | 0.904 | **< 0.001** |

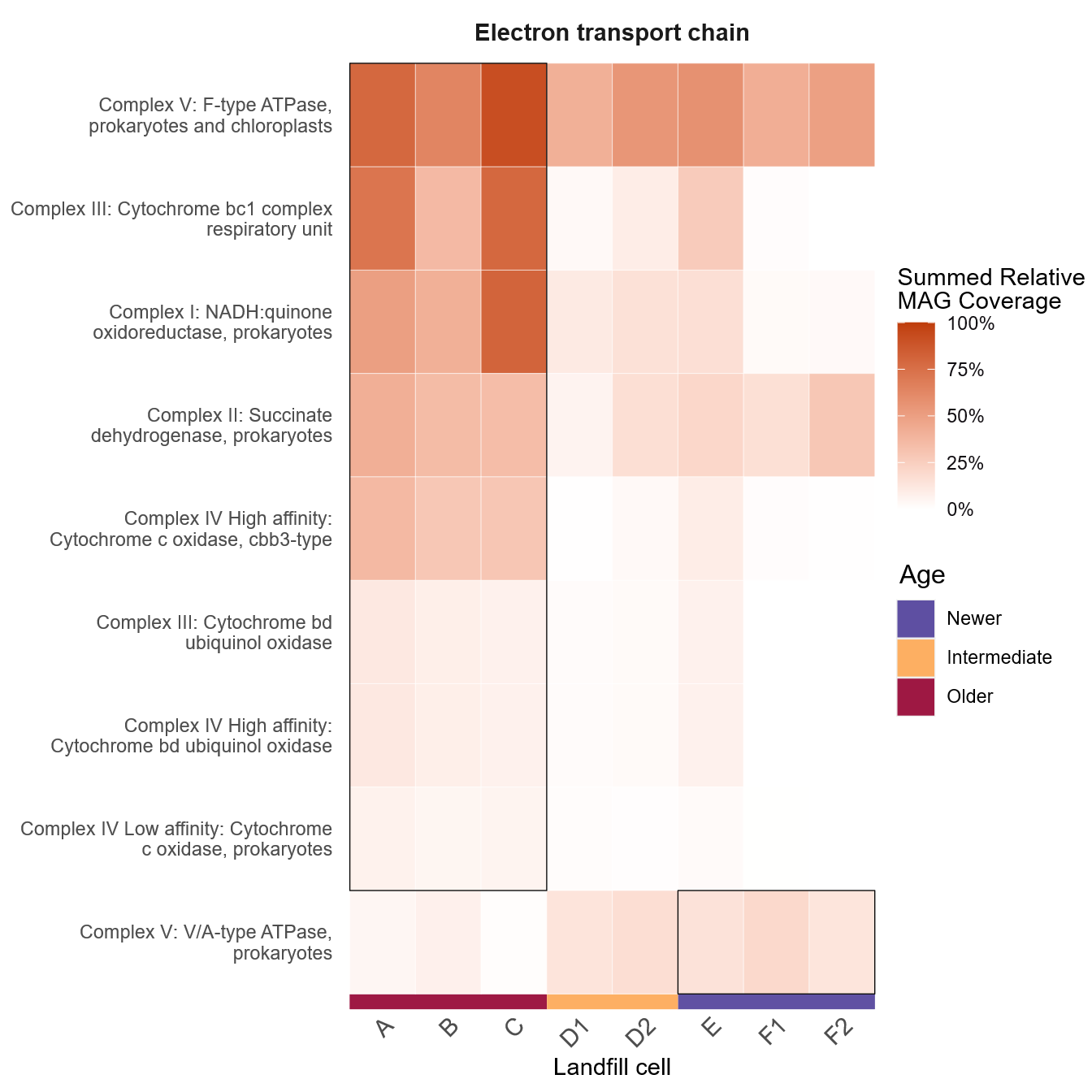

**Figure S1: Electron transport chain component differential abundance across landfill cells.** Heatmap depicting the summed relative coverage of all MAGs in each cell that contain the individual electron transport chain complexes. Complexes were required to be >70% complete within a MAG for inclusion. Only complexes which had significantly different abundances between Older and Newer cells (Wilcoxon rank sum test, Benjamini-Hochberg-corrected p-value of p < 0.05) are displayed. Boxes indicate whether abundance was significantly higher in Older or Newer cells.

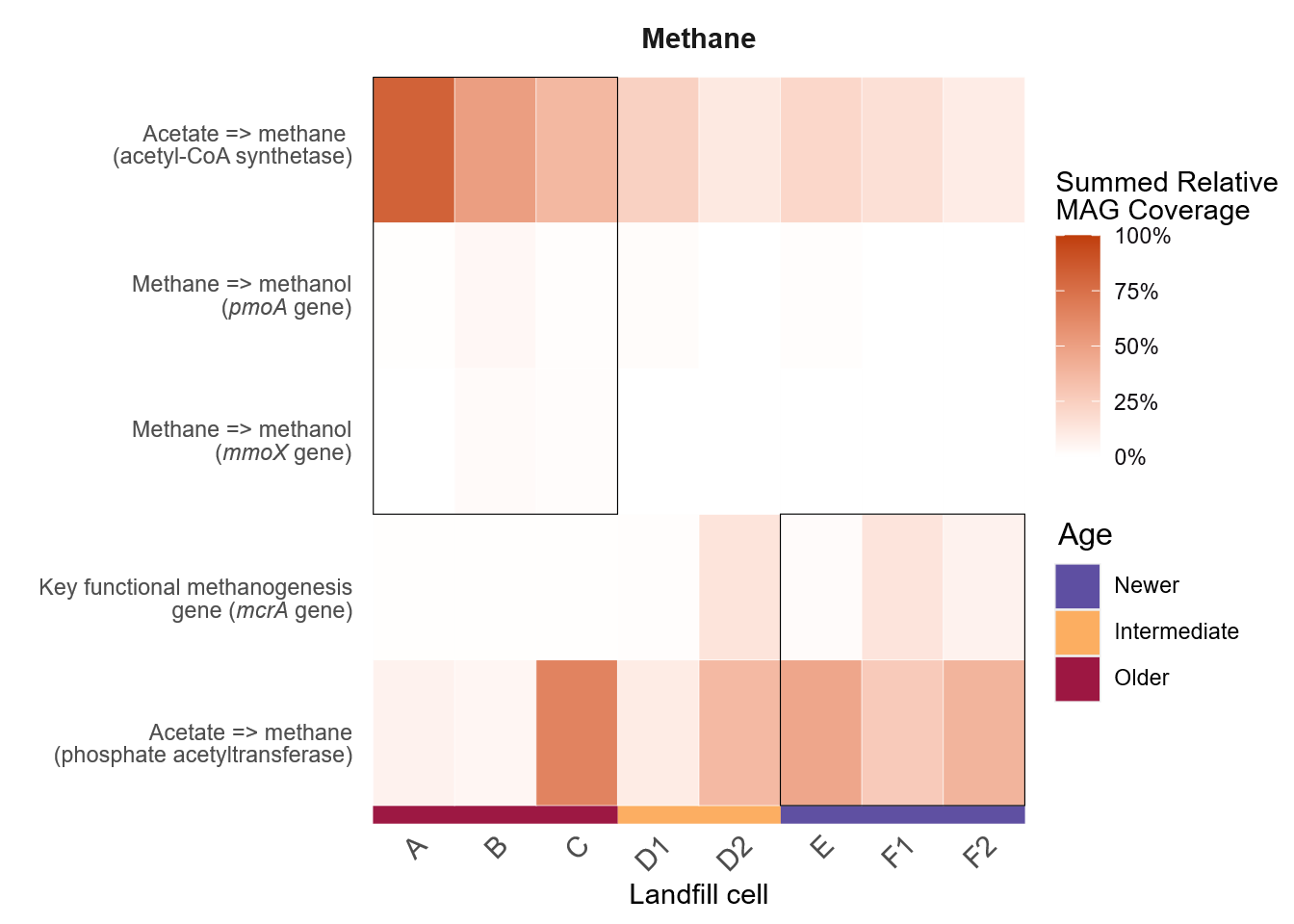

**Figure S2: Methane cycling differential abundance across landfill cells.** Heatmap depicting the summed relative coverage of all MAGs in each cell that contain genes associated with methanogenesis and methanotrophy. Only genes which had significantly different abundances between Older and Newer cells (Wilcoxon rank sum test, Benjamini-Hochberg-corrected p-value of p < 0.05) are displayed. Boxes indicate whether abundance was significantly higher in Older or Newer cells. For all DRAM pathways, the gene(s) of interest are listed in brackets.

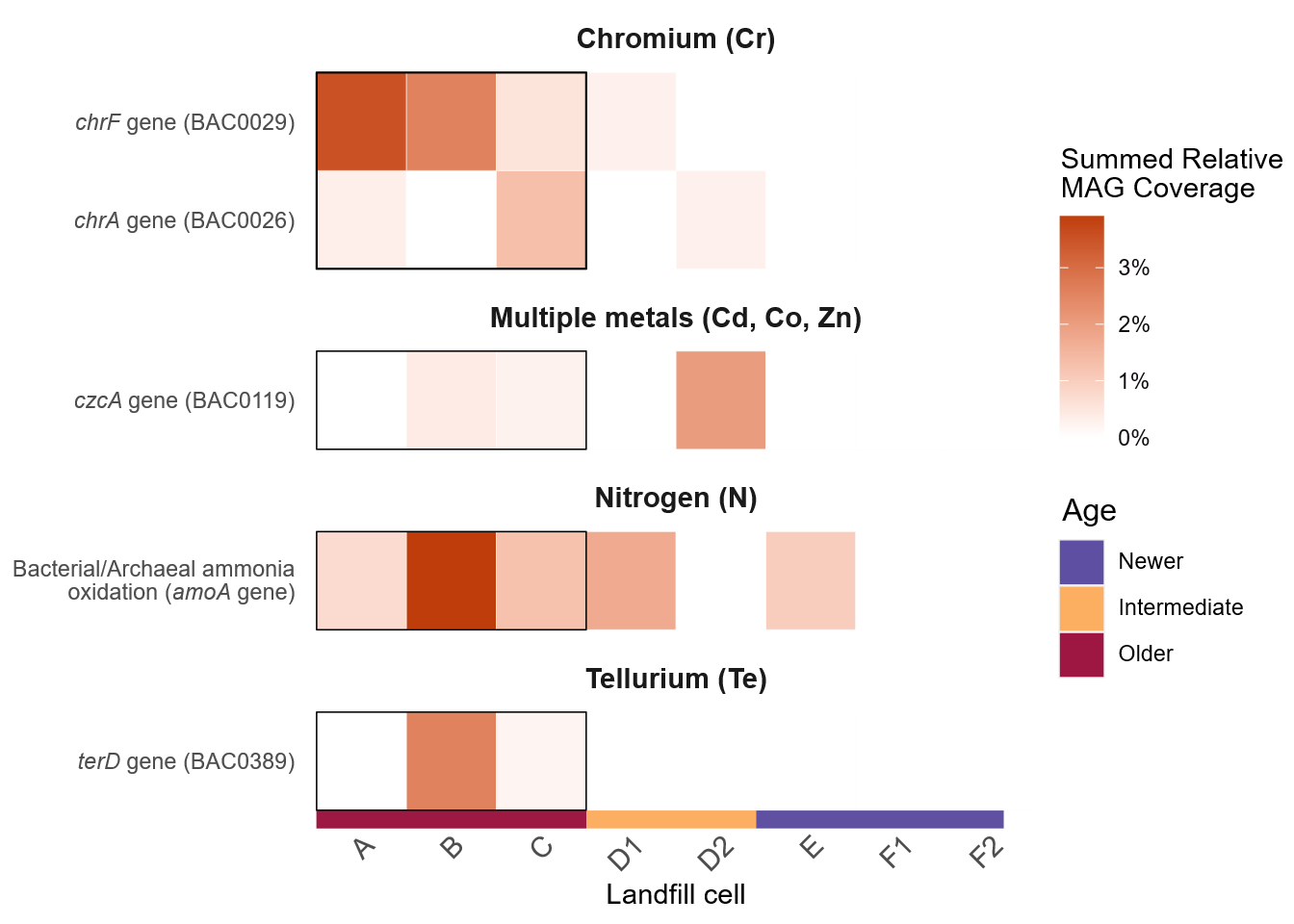

**Figure S3: Abundance across landfill cells of significantly different but low abundance functions.** Heatmap of the summed relative coverage of all MAGs in each cell that contain pathways/genes with a maximum relative coverage below 5%. Only pathways/genes which had significantly different abundance between Older and Newer cells (Wilcoxon rank sum test, Benjamini-Hochberg-corrected p-value of p < 0.05) are displayed. Boxes indicate whether abundance was significantly higher in Older or Newer cells. The processes depicted here were removed from Figures 2 and 3 based on their low abundances. For all DRAM pathways, the gene(s) of interest are listed in brackets. For genes from the BacMet database, the BacMet number is listed in brackets.

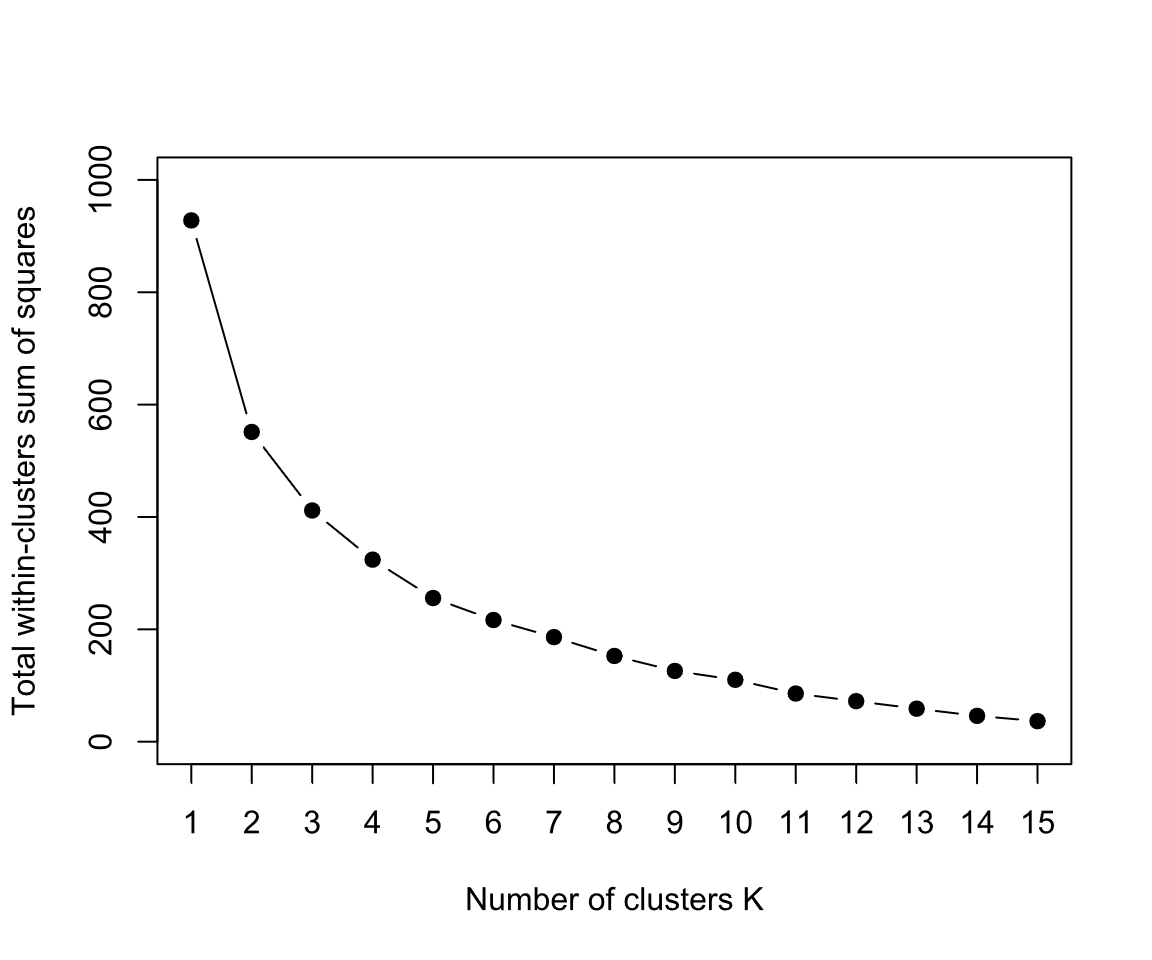

**Figure S4: Elbow plot of within-clusters sum of squares for z-score adjusted 5-year rolling averages of 36 years of landfill leachate chemistry data.**

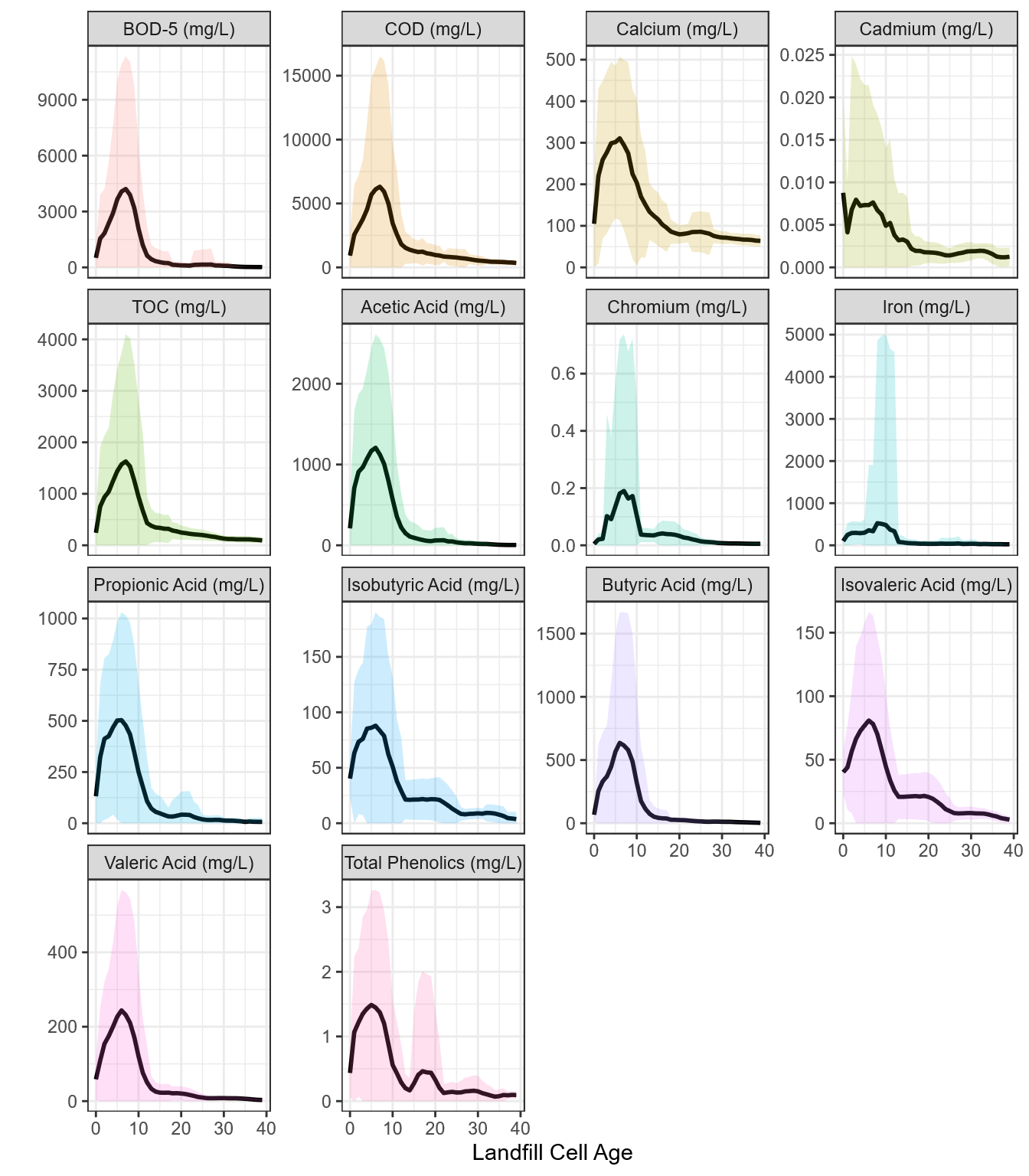

**Figure S5: Line graphs depicting the 5-year rolling average (line) and standard deviation (band around line) for landfill leachate parameters across 36 years of leachate chemistry data.** All depicted parameters were within one k-means cluster (cluster 2, “Organics and metals”).

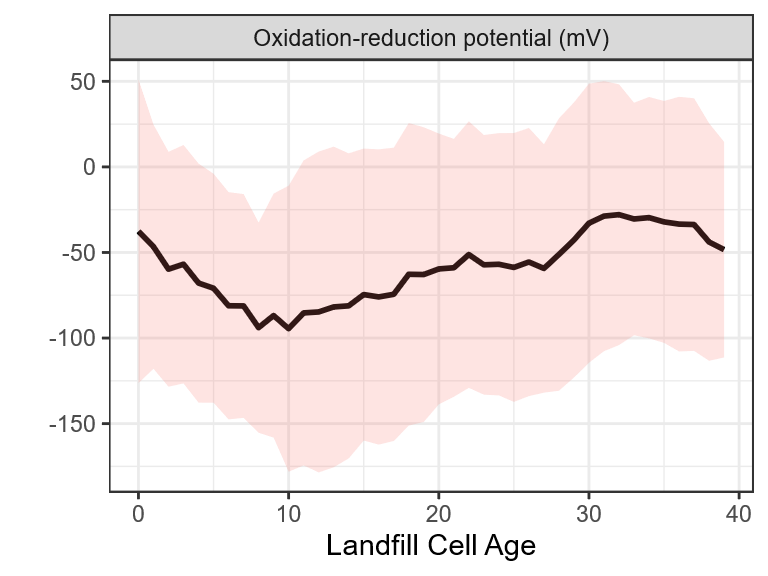

**Figure S6: Line graph depicting the 5-year rolling average (line) and standard deviation (band around line) for oxidation-reduction potential (sole representative of cluster 3, “redox”), across 36 years of leachate chemistry data.**

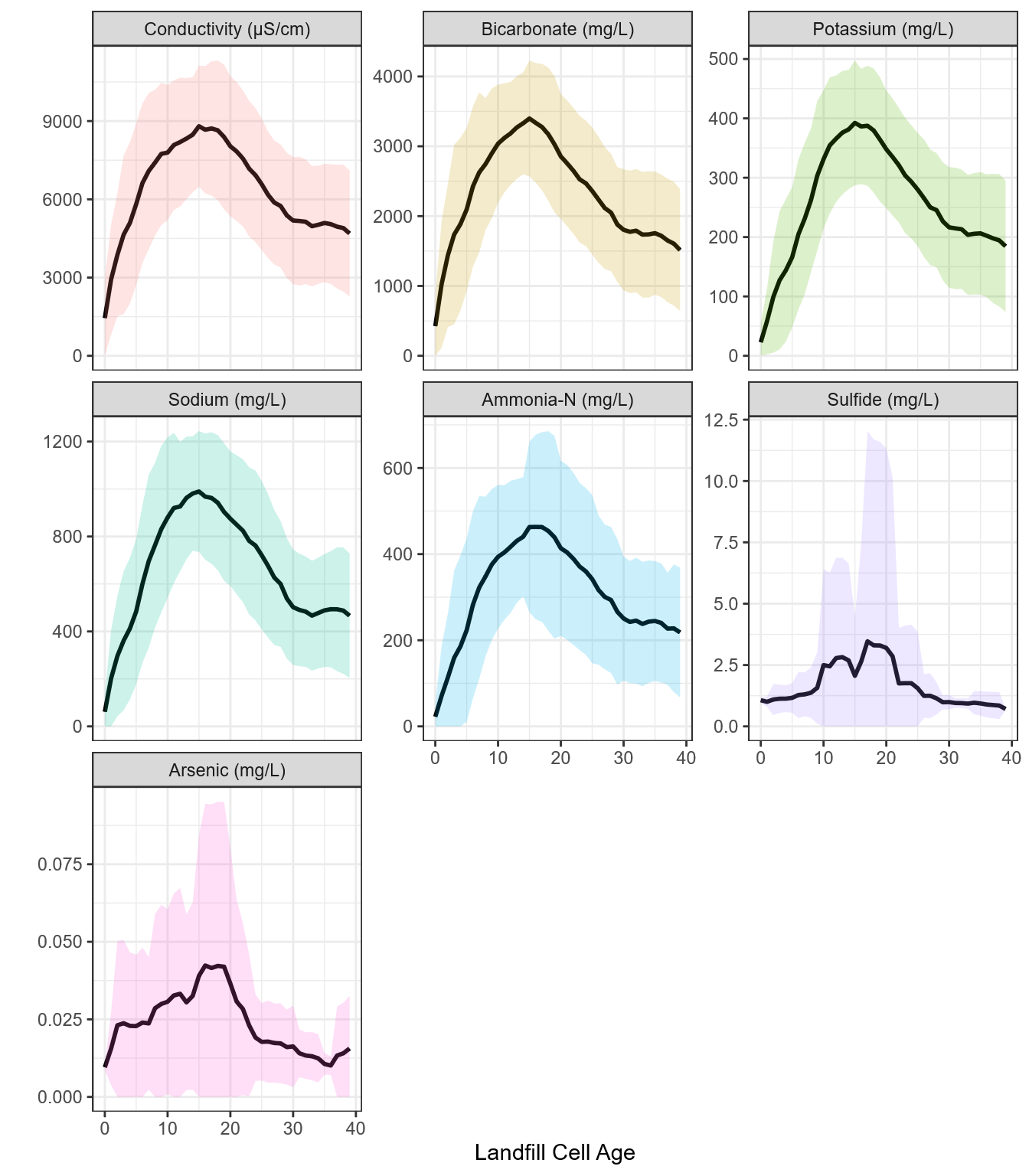

**Figure S7: Line graphs depicting the 5-year rolling average (line) and standard deviation (band around line) for landfill leachate parameters across 36 years of leachate chemistry data.** All depicted parameters were within one k-means cluster (cluster 4, “Conductivity”).

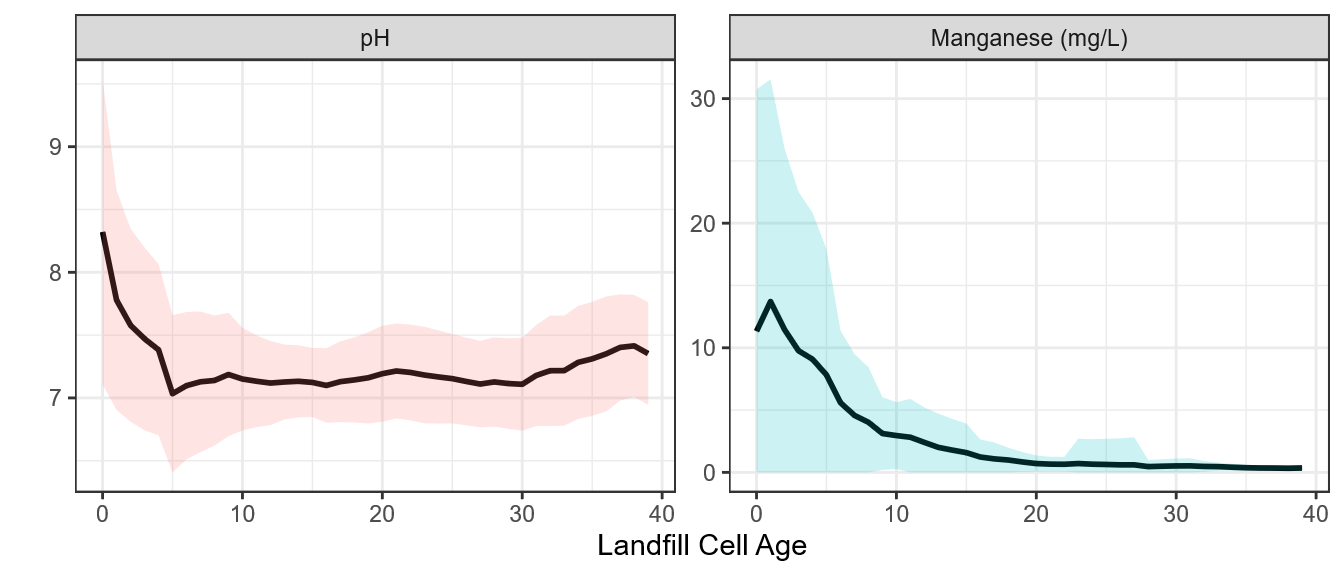

**Figure S8: Line graphs depicting the 5-year rolling average (line) and standard deviation (band around line) for landfill leachate parameters across 36 years of leachate chemistry data.** All depicted parameters were within one k-means cluster (cluster 5, “pH”).

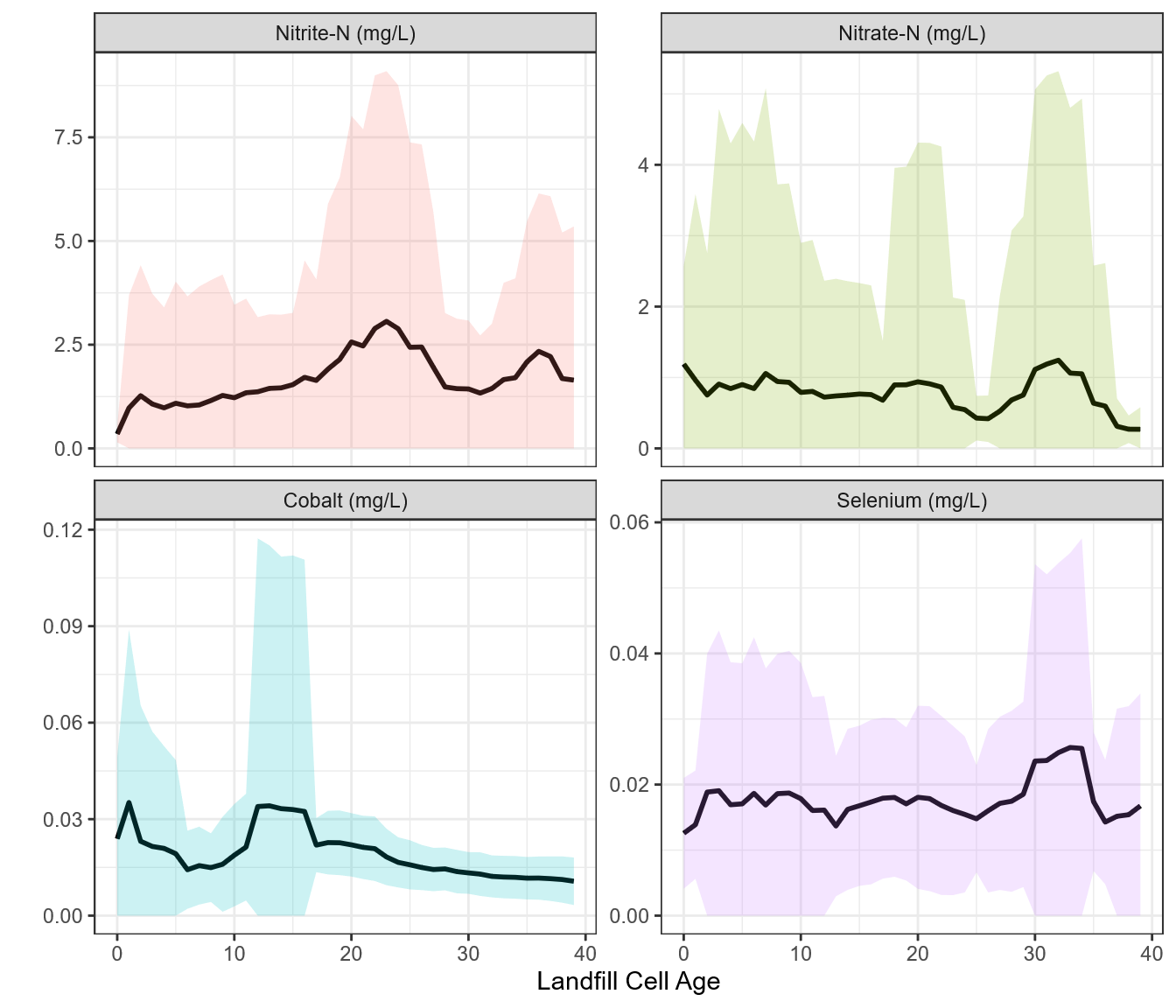

**Figure S9: Line graphs depicting the 5-year rolling average (line) and standard deviation (band around line) for landfill leachate parameters across 36 years of leachate chemistry data.** Depicted parameters were in k-means cluster 6 (nitrite-N), cluster 7 (nitrate-N, selenium), and cluster 8 (cobalt).
